# Organoid-like Functional Adrenal Gland Cortex Derived From Human Pluripotent Stem Cells

**DOI:** 10.64898/2026.06.30.735632

**Authors:** Jessica McAlpine, Christina James, Bhavik Dalal, Kimata Thomas, Trinity Knight, Nadja Zeltner

**Affiliations:** Center for Molecular Medicine, University of Georgia, Athens, GA 30602, USA; Department of Biochemistry and Molecular Biology, University of Georgia, Athens, GA 30602, USA; Department of Cellular Biology, University of Georgia, Athens, GA 30602, USA

**Keywords:** Adrenal gland, adrenal cortex, human pluripotent stem cells, aldosterone, cortisol, DHEA-S

## Abstract

The adrenal cortex is a critical for life endocrine system. It manages metabolic homeostasis, electrolyte balance, stress response, and sex development. This is accomplished through the release of various steroids from a dynamically changing landscape of concentric cellular zones/layers. Adrenocortical dysfunction is implicated in pathologies ranging from adrenal insufficiency to hypertension. The in-depth investigation of adrenal gland biology, pathology and drug discovery has been hampered by a lack of human, experimentally tractable models. Particularly missing are models that recapitulate the cellular diversity of the adrenal cortex with representation of all fetal and adult cell layers and the capsule. Here, we employ human pluripotent stem cells (hPSCs) to generate cells of all three cortex zones alongside capsular cells in a single 2D platform. This platform mimics the cellular diversity expected from an organoid, yet it provides the simplicity of monolayer cultures, that are better suited for drug discovery and high-throughput settings. These cultures secrete zone-specific steroids (cortisol, aldosterone, and DHEA-S) and exhibit robust, physiologically relevant ACTH stimulation responses. Transcriptomic analysis revealed sequential acquisition of profiles consistent with adrenocortical development, zonation, and signaling programs consistent with zone maintenance. Our platform is validated by a literature meta-analysis that defines transcriptomic signatures for each human cortical cell type, spanning fetal and adult stages. Together, this novel adrenocortical platform enables investigation of adrenal development, disease mechanisms, and therapeutic strategies.

**Significance Statement:** We describe a hPSC-based 2D differentiation strategy with the cell type complexity of an organoid and the technical simplicity necessary for high-throughput assays. The platform contains all cortical subtypes that mediate electrolyte regulation, stress response, sex development and self-maintenance of the tissue. This co-differentiation offers a unique opportunity to study human adrenal development, biology, pathology and enables drug discovery.

## Introduction

The adrenal gland is a vital endocrine organ composed of three main components: the medulla, the cortex, and the capsule. The inner medulla mediates the fight-or-flight response through the release of epinephrine (EPI) and norepinephrine (NE) in response to acute stress. The outer cortex regulates body homeostasis, metabolism, and sex development through the secretion of steroid hormones in response to adrenocorticotropic hormone (ACTH), angiotensin II or potassium^1–4^. Surrounding the cortex, the adrenal capsule generates a signaling gradient that maintains cortical zonation and sustains progenitor populations throughout life^1,5,6^.

Proper organization of the adrenal cortex into functionally distinct zones is essential for adrenal function. The outer zona glomerulosa (zG) synthesizes mineralocorticoids (aldosterone) essential for electrolyte regulation. The middle zona fasciculata (zF) produces glucocorticoids (cortisol) that mediates the stress response, metabolism and body homeostasis. Finally, the inner zona reticularis (zR) generates androgen precursor steroids (dehydroepiandrosterone sulfate (DHEA-S), a precursor to estrogen and testosterone^1^, essential for proper sex development. Human adrenal development proceeds through a series of well-defined stages, beginning with the adrenogenic coelomic epithelium (AdCE) at 3-4 weeks post-conception (wpc), which migrate to the developing kidney at around 4-5 wpc and differentiate into adrenocortical progenitors (AdCPs) by 5 wpc^7^. Around this time, medulla progenitors migrate to and subsequently invade the AdCPs, initially appearing as dispersed islands within the fetal cortex and gradually coalescing toward the center of the gland^8^. Shortly thereafter, the human fetal adrenal gland undergoes rapid zonation. By 5-6 wpc, AdCPs differentiate into an outer definitive zone (DZ) and by 6-8 wpc an inner fetal zone (FZ) emerges^7^. Around 6-7 wpc, the adrenal gland is encapsulated by a mesenchymal cell population that expresses GLI1 and RSPO3^7^, establishing a capsule. Throughout life, capsule cells will differentiate into adrenocortical cells of each layer, thus playing a central role in maintenance of the organ, as well as providing the option to adapt to environmental stressors by growing specific zones. The DZ expresses HOPX, MME, and HSD3B2 and retains proliferative, progenitor-like properties, while the FZ expresses CYP17A1, SULT2A1, and CYB5A and specializes in sex hormone precursor synthesis^8,9^. By mid-gestation, a third population of steroidogenic cells, the transitional zone (TZ), emerges at the DZ-FZ interface^8^ and later, at birth, gives rise to the zF. By the third trimester, a subset of DZ progenitors further differentiates into aldosterone-producing cells, giving rise to the zG, which is finalized at birth. Meanwhile, the FZ undergoes apoptosis and involutes, ceasing the production of androgen precursors and therefore does not contribute to the adult adrenal gland^8,10^.

Disruption of zone function or zone organization underlies a wide spectrum of adrenal disorders, which can be life threatening when untreated. Currently, the first line of treatment for primary adrenal insufficiency relies on lifelong hormone replacement therapy^11,12^. This hormone treatment is difficult to dose accurately and cannot respond dynamically to physiological demands, which together can result in morbidity^11^. Additionally, many diseases are caused by or accompanied by adrenocortical dysfunction. For example, cortex-derived aldosterone is needed for immediate survival, and its absence causes salt-wasting and death. Conversely, aldosterone overproduction is a risk factor for hypertension^13–15^, stroke^14^, congestive heart failure^14,16^, and diabetes mellitus^13^. Thus, a simple, experimentally tractable model platform is needed to study human adrenal developmental disorders, adrenal homeostasis, adult pathologies and finally to conduct drug discovery. Current animal models have species specific limitations. For example, human adrenal zonation and structure is different^17–19^, predominant adrenocortical steroids vary^18–20^, and medulla-secreted factors^21,22^ (and subsequently cortex-medulla communication^21^) differ. The availability of untransformed primary adrenocortical tissue from humans is scarce, precluding high-throughput physiological and pathological investigations.

The recent availability of human pluripotent stem cell (hPSC) technology has revolutionized the study of human development and diseases^23,24^. Sakata *et al.* established a directed differentiation protocol for FZ-like adrenocortical cells from hPSCs through modulation of NOTCH, ACTIVIN, and WNT signaling^25^. Building on this, Mayama *et al.* generated adrenocortical organoids capable of cortisol synthesis and possessing a DZ. However, their capsule-like RSPO3-expressing cells were differentiated separately and assembled with cortical progenitors and did not emerge endogenously within the organoids^5^. Additionally, Mayama et al. did not report aldosterone synthesis, suggesting that a signaling component may still be missing from their system^5^. Thus, a hPSC-based adrenocortical model system that contains all zones and the capsule and functionally releases all steroids in a stimulus responsive manner is currently lacking.

Here, we establish a platform for generating adrenocortical cells that have the capacity to synthesize and secrete steroids from all three adrenocortical zones, i.e. cortisol, DHEA and aldosterone, from hPSCs. They recapitulate canonical stages of human adrenocortical development and differentiate alongside capsule cells in a single culture system. Given the diversity of cell types and functions in this system, it is comparable to an organoid, yet it is grown in a 2D platform, providing the advantage of simplicity and adaptation to high-throughput analysis. We further show that these adrenocortical cells respond to ACTH in a manner comparable to human primary fetal adrenocortical cells. Furthermore, in a meta-analysis of nine recent human adrenocortical sequencing papers, we established a consensus gene signature for each human adrenocortical cell type and the capsule, including previously poorly defined populations such as zG progenitor cells. We used these expression signatures to identify cell types from all cortex zones in our cultures. Together, we provide a powerful new tool to investigate human adrenal gland development, pathology and enable drug discovery.

## Results

### Adrenocortical Progenitor Generation

To develop a differentiation paradigm for AdCPs, we modified an approach that generates gonad progenitor cells, the adrenal cortex’s closest developmental relative^26^ (**Supp. Fig. 1A**). Lineage tracing studies in mouse identified a shared adrenogonadal primordium expressing Nr5a1 and Gata4^7,27^. However, single-cell RNA sequencing (scRNAseq) studies on human fetal adrenals have revealed important species-specific differences between mouse and human adrenocortical development^1,7,28–30^. Human AdCPs do not share a common precursor pool with gonad progenitors, nor do they express GATA4^7^. Thus, we use NR5A1 as a positive marker, alongside GATA4 as a negative marker for AdCP differentiation. As adrenal precursors develop medial to gonad precursors^7,26^ and BMP4 is lateralizing^15^, we titrated BMP4. Furthermore, as FGF9 is known to induce kidney and gonad progenitor gene expression^25,26,31^, we titrated FGF9 alongside BMP4. We found that removing BMP4 completely and decreasing FGF9 to 20 ng/mL, yielded strongest *NR5A1* expression (RT-qPCR at day 8), coupled with a significant decrease in *GATA4* expression compared to gonad^26^ and kidney^31^ differentiations (**Fig. Supp. 1B**). By immunofluorescence (IF), we confirmed a modest increase in AdCP differentiation efficiency compared to gonad based on the increase of cells positive for NR5A1 and negative for GATA4 (**Fig. Supp. 1C**). In the intermediate mesoderm, anterior-posterior specification is determined largely by the duration and magnitude of WNT signaling, whereby shorter and/or weaker WNT signals generally specify towards anterior and vice versa for posterior fates^31^. Advances in our understanding of human adrenal development have highlighted that adrenocortical specification occurs anterior to the gonad^7^. Considering these discoveries, anterior-posterior specification was optimized by modifying the duration and concentration of the GSK3b-inhibitor Chir99021 (Chir, **Fig. Supp. 1D**). To narrow down a range of time and concentration combinations for deeper investigation, we performed a mini screen by IF. Chir was added for either 2, 3, 4, or 5 days at concentrations of 2-4.5µM in 0.5µM increments. The most apparent NR5A1 increase was observed at days 2 and 3 at concentrations between 3-4.5 µM. (**Fig. Supp. 1E**). Further testing by RT-qPCR in these conditions revealed a significant increase in *NR5A1* without a notable increase in *GATA4* expression when treating with 4µM Chir for 3 days (3d4µM) compared to the original treatment, 3µM for 4 days (4d3µM, **Fig. Supp. 1F**). By IF quantification, we found a 5-fold increase in AdCP differentiation efficiency at day 8 (**Fig. Supp. 1G**) and confirmed that increased *NR5A1* expression was consistent over time, with significantly higher expression at days 8, 12, and 15 (**Fig. Supp. 1H)**. To verify that this change in Chir concentration and duration improved efficiency through anterior-posterior patterning, RT-qPCR was performed for the anterior adrenal HOX gene *HOXB4*, and the posterior gonad HOX gene *HOXD13*^7^. While the decrease in *HOXD13* expression did not have statistical significance (p= 0.08), *HOXB4* expression was significantly increased (p = 0.005, **Fig. Supp. 1I**). To support reproducibility, phase contrast images taken every 2 days of AdCP differentiation are provided (**Fig. Supp. 1J**). By starting with a gonad differentiation protocol, decreasing FGF9, removing BMP4, and adjusting the concentration and duration of WNT signaling, we established AdCP differentiation protocol.

### Characterization of Adrenocortical Progenitor Differentiation over Time

To further validate the identity of our AdCPs, we next established that they pass through appropriate developmental milestones by characterizing their differentiation over time using markers of each developmental stage. Adrenocortical development begins in the early posterior primitive streak (PS; Brachyury/T) that gives rise to a portion of the mesoderm between the lateral plate mesoderm and the paraxial mesoderm, called the intermediate mesoderm (IM). Within the IM, lineages are defined along an anterior-posterior axis, with the early posterior IM (OSR1+/T-) giving rise to the future adrenocortical and gonad cells. The future adrenal cortex is specifically defined once the early/pre-migratory adrenal coelomic epithelium (early AdCE; WT1, GATA5, KRT8) differentiates. This differentiation and migration is followed by upregulation of late AdCE markers (DLK1, NR0B1, NR5A1) and finally formation of AdCPs (NR5A1, OSR2, CITED2, STAR; **Fig. 1A**)^7^. In our directed AdCP differentiation protocol (**Fig. 1B**), PS induction occurs between days 0-2, indicated by upregulation of *T*. Between days 2-4, *T* expression sharply decreases, followed by upregulation and peak of IM marker expression (*OSR1*, **Fig. 1C**, **D**). To test for potential early contamination via off-target differentiation, we performed RT-qPCR and found that markers of endoderm (*SOX17*), non-neural ectoderm (*TFAP2A*), paraxial mesoderm (*TBX6*) were weakly upregulated, while *FOXF1* expression was stronger, indicating a somewhat lateral plate-like identity (**Fig. 1E**). Following IM induction, early AdCE gene expression is upregulated between days 4-6, followed by late AdCE expression increasing between days 6-8 (**Fig. 1F**). Finally, AdCP marker expression emerges by day 8 and peaks around days 12-15 (**Fig. 1F**). Expression of AdCP markers (CITED2, NR0B1, NR5A1) without pervasive GATA4 expression at days 6, 8 and 10 was confirmed with immunofluorescence (**Fig. 1G)**. This sequential expression of developmental milestone markers indicates the emergence of true AdCP cells. *NR5A1* expression is significantly upregulated in the AdCP protocol compared to gonad (RT-qPCR, **p = 0.008; **Fig. 1H**). At day 8, AdCP differentiation efficiency averages approximately 60% compared to around 1% using the gonad differentiation protocol (**Fig. 1I**, IF; **Fig. 1J**, image quantification). OSR2 positive cells were WT1 negative, further supporting AdCP identity of these cells (**Fig. 1K**). Expression of gonad and kidney markers (*LHX9*, *LHX1,* respectively) are low at day 8, indicating minimal IM-derived off-target contamination (**Fig. 1L**). The robust upregulation of adrenocortical-specific marker expression achieved with this protocol is evidence that these cells have the potential to differentiate further into adrenocortical lineages.

**Fig. 1:**
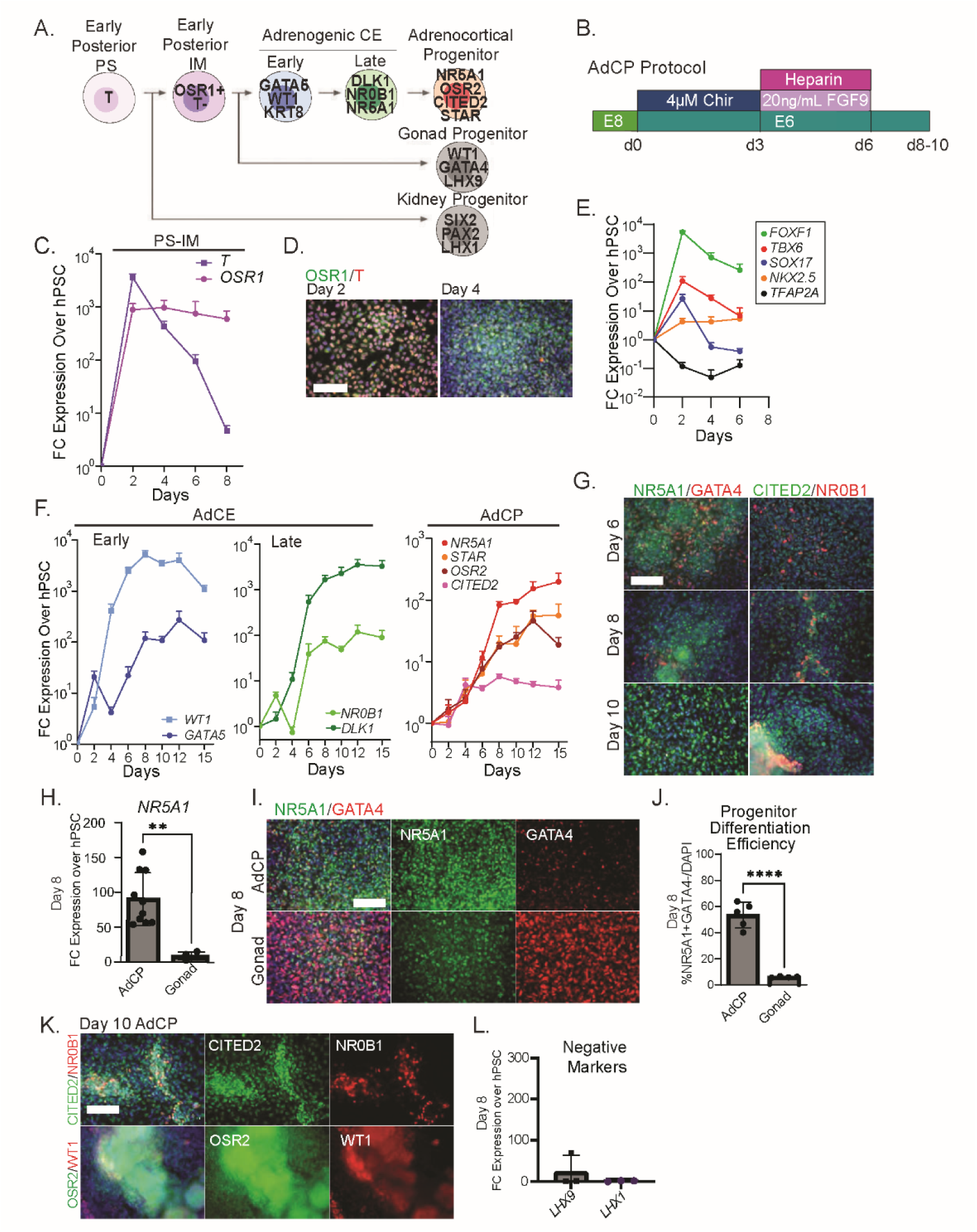
Adrenocortical Progenitor Cell Differentiation Protocol Characterization. (**A**) Diagram of embryonic AdCP differentiation and their respective markers. (**B**) Diagram of AdCP differentiation protocol. (**C**) RT-qPCR for PS and IM markers over time (d2, n= 2-3; d4, n= 4-6; d6, n=9-10). (**D**) IF for posterior IM markers OSR1, Brachyury (T) on days 2 and 4. Scale = 100µm. (**E**) RT-qPCR for early developmental off-target cell type markers from days 0 to 6 (n=3-5). (**F**) RT-qPCR for early AdCE (*WT1*, *GATA5*), late AdCE (*NR0B1*, *DLK1*), and AdCP (*NR5A1*, *STAR*, *OSR2*, *CITED2*). d2, n= 2-3; d4, n= 4-6; d6, n=9-10; d8, n=7-16; d10-d15, n=5-9 (**G**) IF for early cortex development markers (NR5A1, OSR2, CITED2) and gonad progenitor marker GATA4 over time. Scale = 100µm. (**H**) Expression of *NR5A1* and *GATA4* by RT-qPCR in AdCP and gonad differentiation protocols at day 8. Unpaired t-test (n=4-10, **p = 0.008). (**I**) IF for NR5A1 and GATA4 in AdCP and gonad differentiation protocols at day 8 and nuclear counterstain DAPI (blue). Scale = 100µm. (**J**) AdCP differentiation efficiency by quantification of NR5A1+/GATA4- cells. Unpaired t-test (n = 4; ****p < 0.0001). (**K**) IF of day 10 AdCP cells. Nuclear counterstain DAPI (blue). Scale = 100µm. (**L**) Expression of markers for developmentally related off-target cell types (*LHX1*, *LHX9*) at day 8 by RT-qPCR. All graphs in this figure show mean ± SEM. Abbreviations: AdCE = adrenogenic coelomic epithelium; AdCP = adrenocortical progenitor; IM = intermediate mesoderm; IF = Immunofluorescence; PS = primitive streak.

### TGF-b Inhibition is Crucial for AdCP to Definitive Zone Transition

AdCPs have been reported to differentiate either into the DZ or directly into cells of the FZ^7^. The embryonic DZ is a progenitor pool for the steroidogenic cortex through the second trimester marked by expression of HOPX, HSD3B2, NOV/CCN3, and MME^7^. Aldosterone production begins in the DZ, creating a heterogeneous cell population comprised of both aldosterone producing cells and non-steroidogenic DZ progenitors (**Fig. 2A**). Thus, to generate diverse steroidogenic cell types, we aimed to guide AdCPs through this DZ intermediate. As TGF-b is known to diminish adrenocortical survival *in vitro*,^32^, and B-27 supplement supports cholesterol metabolism and related processes^33^, AdCPs were treated with the TGF-b inhibitor SB431542 (SB) and B-27 supplement (B27) to encourage differentiation towards DZ. SB and B27 treatment began either at day 6, immediately following FGF9 treatment (d6DZ), or at day 10, once AdCP identity is established (d10DZ, **Fig. Supp. 2A**). By day 15, expression of AdCP markers (*NR5A1*, *OSR2*) were higher in the d10DZ condition than in d6DZ or in E6 alone. Additionally, *WT1* expression was higher in d6DZ than in d10DZ or E6 at day 15, suggesting that starting the TGF-b inhibition before cells have established an AdCP identity may induce off-target differentiation (**Fig. Supp. 2B**). Expression of DZ markers (*HSD3B2*, *NOV*, *HOPX*, and *MME*) was evaluated on days 15, 22, and 32 in all 3 conditions (**Fig. Supp. 2C**). d10DZ expressed significantly more *NOV* and *HSD3B2* at days 22 and 32, while HOPX and MME was upregulated in all three conditions (**Supp. Fig. 2C**). These differences were confirmed at the protein level (**Supp. Fig. 2D**). Co-expression of NR5A1 and MME was demonstrated in the d10DZ condition (**Fig. Supp. 2E**). To evaluate the maintenance of proliferating NR5A1+ cells over time, IF was performed for KI67 and NR5A1 in each condition. As many of the NR5A1+ cells were indeed cycling (KI67+, **Fig. Supp. 2F**), this adds support for their progenitor-like identity. We therefore selected d10DZ as our DZ differentiation condition. Going forward, we will call these cells DZ cells (**Fig. 2B**).

**Fig. 2:**
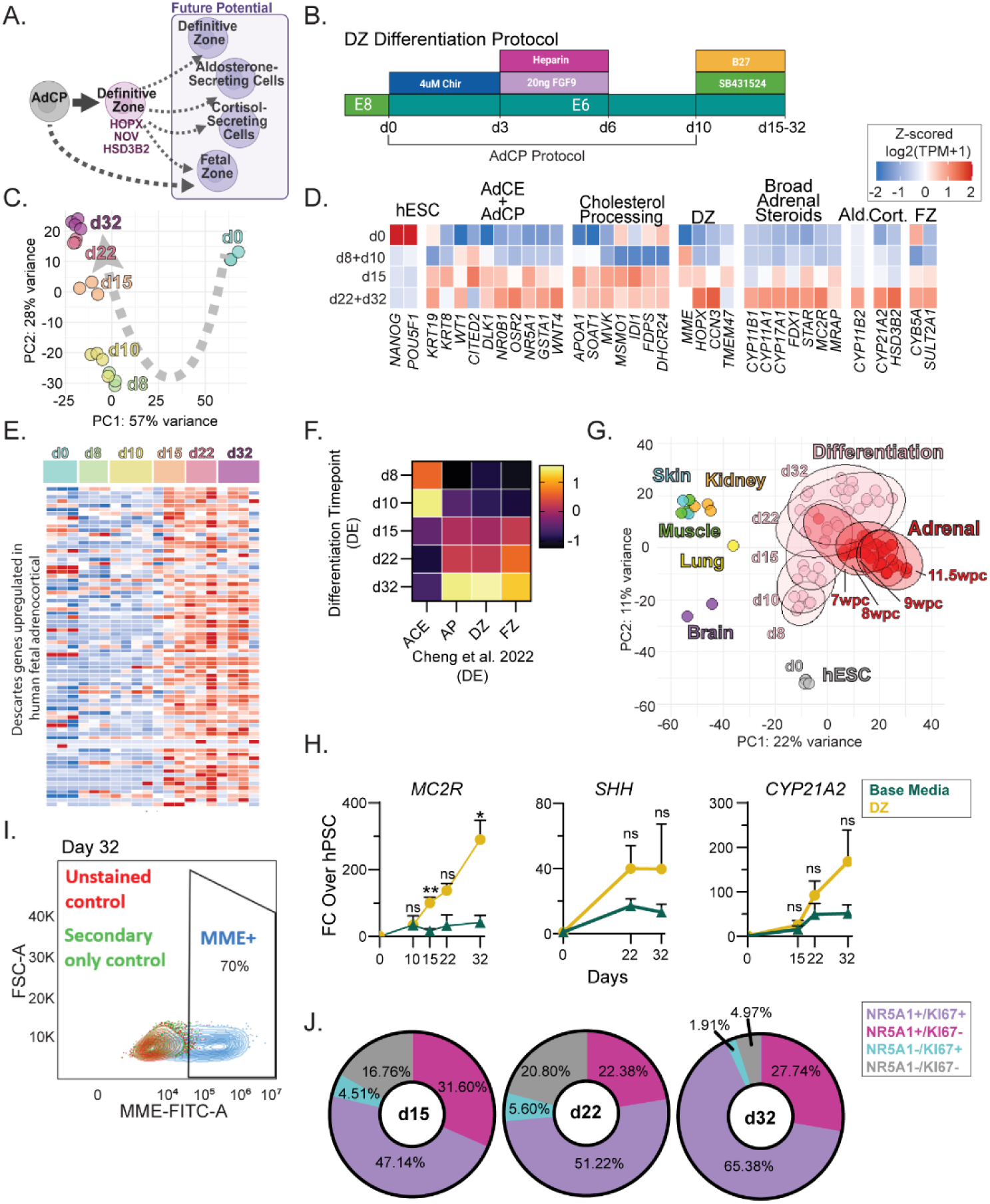
Differentiation of Adrenocortical Progenitors into Definitive Zone Cells. (**A**) Diagram depicting the differentiation potential of AdCP and DZ cells during adrenocortical development. (**B**) Diagram of differentiation method for DZ cells. (**C**) PCA plot of DZ differentiation over time characterized by bulk RNAseq. (**D**) Heatmaps of relative expression of canonical adrenocortical development and function markers and (**E**) computationally identified fetal adrenocortical markers from the Descartes database^29^ over time; expression = z-scored log2(TPM+1). (**F**) Heatmap of correlation between transcriptional profiles at each DZ differentiation timepoint and markers of adrenocortical developmental cell subtypes identified in Cheng *et al*. 2022^7^. (**G**) PCA plot comparing transcriptional profiles of differentiated DZ cells and multiple fetal tissues from del Valle *et al.* 2023^1^. (**H**) RT-qPCR for ACTH receptor *MC2R* (n = 3-6) and zonation maintenance factor *SHH* (n = 3-4) over time in DZ compared to E6. Steroidogenic potential was assessed with RT-qPCR for *CYP21A2* in E6 and DZ conditions over time (n=3-6). Multiple unpaired t tests with Holm-Šídák multiple comparisons corrections (*MC2R*: d15, **p = 0.003; d32, *p = 0.049). (**I**) FACS of day 32 DZ cells for the DZ marker MME. (**J**) Quantification of IF days 15, 22, and 32 for NR5A1 and the proliferation marker KI67 to identify putative cortex progenitor cells of relative proportions of populations positive for each marker. All graphs in this figure show mean ± SEM. Abbreviations: AdCP = adrenocortical progenitor; DZ = definitive zone; E6 = base media; IF = immunofluorescence; PCA = principal component analysis; TPM = transcripts per million.

### Characterization of Definitive Zone Cells

DZ cells’ transcriptional profiles were analyzed by bulk RNAseq. from pluripotency (day 0) through day 32 (**Fig. 2C, Fig. Supp. 3A**). By PCA, we observed that our DZ cells followed a continuous trajectory of differentiation in order by time, indicating a progressive temporal maturation. Days 8 and 10 clustered together by PCA, as did days 22 and 32, indicating that these clusters may reflect distinct stages in differentiation. (**Fig. 2C**). We next examined whether these coordinated transcriptional changes over time reflected adrenocortical identity across each distinct stage of differentiation. Testing for markers of adrenocortical specification revealed a peak in expression of cholesterol processing, steroidogenic, and DZ markers by late differentiation on days 22 and 32 (**Fig. 2D**, **Fig. Supp. 3B**). Additionally, differentiating DZ cells were tested for markers enriched in fetal adrenocortical cells from the Descartes cell type database^29^. We found a gradual increase in expression of adrenocortical genes from day 15 onwards (**Fig. 2E**). Comparison of DZ cells to human fetal cortex cell types described in Cheng et al. 2022^7^ by bulk RNAseq using correlation analysis revealed that differentiating cortex progenitors are transcriptionally more AdCE-like through days 8 and 10 in agreement with our RT-qPCR results in **Fig. 1F**. They only adopt a more AdCP-like profile by day 15. The strongest correlation to embryonic adrenal gland tissue showed up by day 32 and revealed that day 8 and 10 cells most closely resembled AdCE, while from d15 onwards there was a gradual increase in the overlap in the expression of AdCP, DZ, and FZ markers (**Fig. 2F**). Our data sets were also compared to publicly available RNAseq datasets of fetal human whole embryo tissues and showed the closest similarity to the fetal adrenal (ArrayExpress/Biostudies accession number E-MTAB-12492^1^, **Fig. 2G**). This indicated that our older cultures most closely resemble 7 wpc in humans (**Fig. 2G**), however, data shown later (**Fig. 5**) shows that our cultures are even more mature than that. Importantly, DZ cells expressed *MC2R* and *SHH*, indicating the potential for signaling to capsule cells^6,34,35^ and for responding to ACTH (**Fig. 2H**). Expression of MME/CD10 was also evaluated by FACS, with 70% of cells expressing MME by day 32 (**Fig. 2I**). Quantification of proliferating cortex cells (NR5A1+/KI67+) identified by IF revealed that nearly one third of the cells were proliferative NR5A1+ progenitor cells by day 15, around 22% by day 22, and 27% by day 32, indicating that DZ cultures contain a relatively stable population of cycling cortex cells over time (**Fig. 2J**; **Fig. Supp. 2F**). Steroidogenesis markers increased gradually over time at the transcript level until day 32 both by bulk RNAseq (**Fig. 2D**, **Supp. Fig. 3B**) and RT-qPCR (**Fig. Supp. 3C**). Together, these results indicate that the DZ differentiation protocol results in a robust induction of DZ cells by day 15 and onward and that a subset of these DZ cells may be poised to become steroidogenic cells.

### Continuous DZ Media Treatment Supports Steroidogenic Cell Differentiation

While it is known that DZ cells differentiate into cortisol-secreting cells, it is unknown whether AdCPs can differentiate into cortisol-secreting cells directly, as is the case with FZ cells^7^, or whether a DZ intermediate is required. Thus, to identify the best approach to differentiate cortisol-secreting cells, we utilized both a direct and a stepwise approach. For the direct approach, AdCPs were treated with steroidogenesis induction media containing low ACTH (8.8pM), IGF2 (25ng/mL), and CRH (40pM)^36^ (AIC media), consistent with circulating basal ACTH concentrations during fetal development. For the stepwise approach, AdCPs were first treated either with base media (E6) or DZ media until day 15. On day 15, E6 or DZ media either continued or was changed to AIC. For all conditions, steroidogenic potential was assessed by stimulation with AIC media containing a strong pulse of 10nM ACTH for 48 hours (**Fig. Supp. 4A**). Each condition was evaluated for expression of the enzymes required for cortisol synthesis, CYP11B1 and CYP21A2 by IF. By day 32, we observed expression of both markers in cells treated only with DZ media (DZ→DZ, **Fig. Supp. 4B**). A similar trend was observed by RT-qPCR, which demonstrated that *CYP21A2*, *HSD3B2*, and *CYP11B1* expression were highest in the DZ→DZ condition. However, DZ→AIC treatment led to similar expression levels of *CYP11B1* and *SULT2A1,* while lacking robust *HSD3B2* expression. Thus, the DZ→AIC condition appears to represent a more FZ-like phenotype (**Supp. Fig. 4C**). Cortisol ELISA revealed that DZ→DZ cells secreted more cortisol than other conditions by day 32 (**Fig. Supp. 4D**), in line with evidence of steroidogenic identity suggested by its gene expression profile. Furthermore, only the DZ→DZ treatment resulted in aldosterone production (**Fig. Supp. 4E),** whereas both DZ◊DZ and DZ◊AIC conditions led to DHEA-S secretion (**Fig. Supp. 4F**). Altogether, these results indicate that continuous treatment with AIC prevents cortisol-and aldosterone-secreting cell identity while promoting a more FZ-like identity. Furthermore, they indicate that the DZ→DZ condition produces all three steroids cortisol, DHEA and aldosterone. To our knowledge, this is the first report thereof. Going forward we will call this matured DZ (mDZ).

### Matured DZ Cells Respond Robustly to Stimulation with ACTH

The capacity for responding to ACTH stimulation is an important quality of an adrenocortical model system. ACTH acts as a stimulus for steroid release in response to stress through the coordination of its receptor MC2R and its accessory protein MRAP. Thus, we next assessed the mDZ cells’ response to ACTH. Different adrenocortical cells are specialized in the type of steroids they produce, based on which specific steroidogenic enzymes they express (**Fig. 3A**). We found that without stimulation, mDZ cells are competent to secrete steroids of all 3 cortex zones by day 32 (**Fig 3B**). As we had already established, that they express the ACTH receptor *MC2R* (**Fig. 2H**), we next tested whether the secretion of steroids would increase in response to ACTH. Comparing the cortisol output of stimulated and unstimulated mDZ cells at d32 reveals a robust ACTH response by day 32 with a 16-fold increase in cortisol release upon stimulation (**Fig. 3C**). DHEA-S release increased similarly upon stimulation (**Fig. 3D**). Additionally, we found that the stimulation-dependent cortisol release occurs in an ACTH dose-dependent fashion, confirming that cortisol secretion is driven specifically by ACTH-MC2R signaling (**Fig. Supp. 5A**). Doses above 100nM ACTH yielded lower extracellular concentrations of cortisol after 48 hours, possibly indicating ER stress (**Fig. Supp. 5A**). This cAMP signaling-induced cortisol secretion was confirmed with 8-br-cAMP in a dose dependent fashion, providing orthogonal validation for their steroidogenic response to PKA signaling (**Fig. Supp. 5B**). These adrenocortical cells express major steroidogenic enzymes at the protein level (HSD3B2, FDX1, CYP21A2, CYP11B1, CYP11A1, **Fig. 3E**). These markers were co-expressed with NR5A1 as expected. Furthermore, they bore characteristic steroidogenic cortex cell morphology with polyhedral cell bodies and round nuclei, further validating their adrenocortical identity (IF, **Fig. 3E**). Steroidogenic functional identity was further confirmed with transmission electron microscopy (TEM) to characterize cellular ultrastructure, revealing mitochondria near lipid droplets, which are characteristic of adrenocortical cells cultured *in vitro*^37^ (**Fig. 3F)**. Together, these results reveal that the mDZ cells functionally recapitulate important qualities of human adrenal cortex cells.

**Fig. 3:**
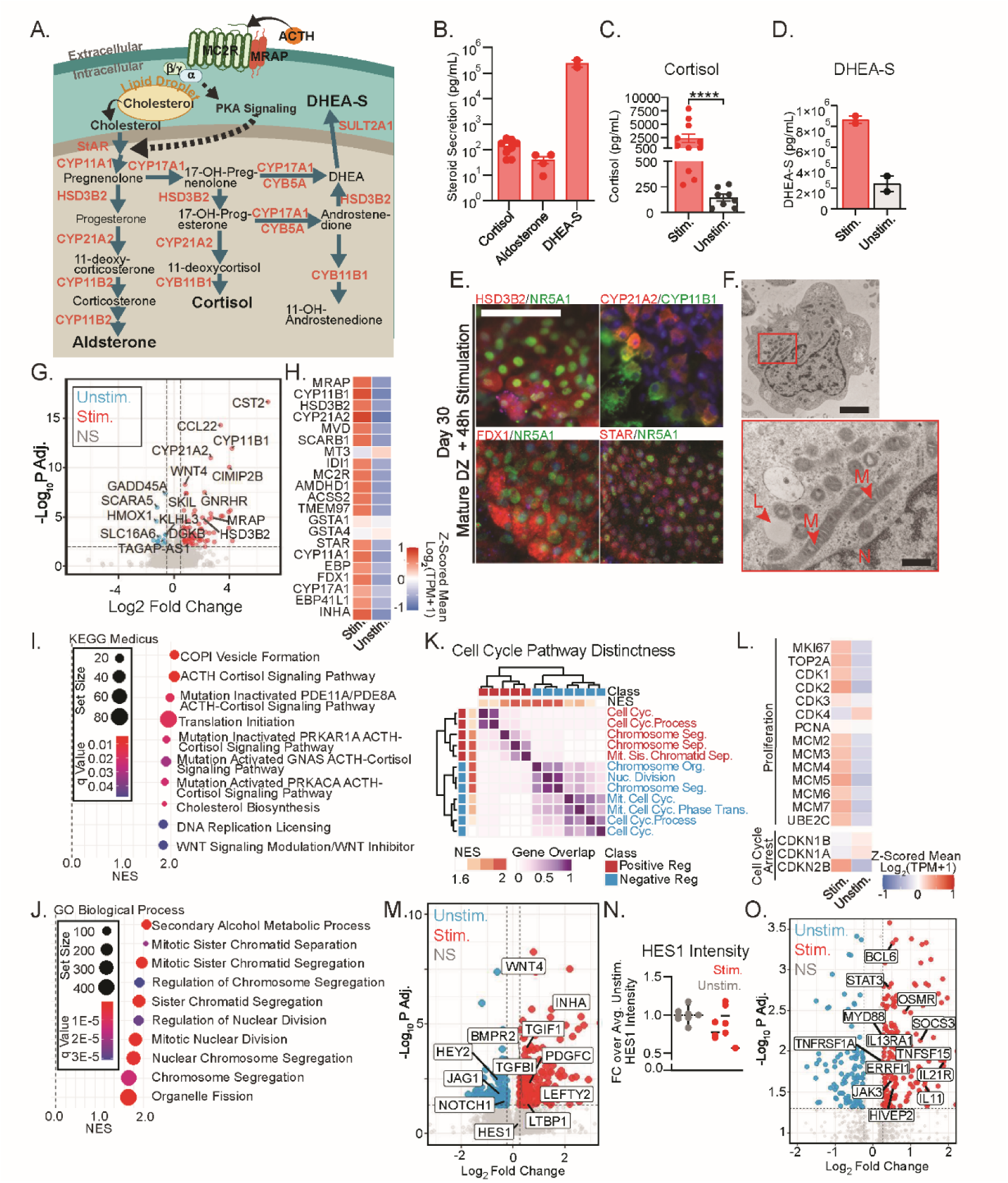
Steroidogenic Induction and Functional Validation of Matured DZ Cells. (**A**) Diagram of adrenocortical steroidogenesis pathways. (**B**) Secretion of steroids at differentiation day 32 without stimulation (ELISA). (**C**) Cortisol release from day 32 mDZ upon 48-hour stimulation with AIC versus unstimulated. Statistics by Mann-Whitney test (****p < 0.0001, n = 8-11). (**D**) DHEA-S release from mDZ cells upon stimulation compared to without stimulation (ELISA; n = 2). (**E**) IF of AIC-stimulated mDZ cells at day 32 detecting steroidogenesis markers, counterstained with DAPI; 100µm scale. (**F**) TEM of day 32 mDZ (top; 2µm scale); magnification of region indicated in red box (bottom; 400nm scale). Abbreviations: N, nucleus; L, lipid droplet; M, mitochondria. (**G**) Volcano plot of top 15 differentially expressed genes in stimulated mDZ (right) compared to unstimulated (left). Genes considered not significant (NS) when log_2_ fold change <=|0.5| (x-axis) or adjusted p-value <= 0.05 (y-axis). (**H**) Heatmap of relative expression of transcripts reported to be upregulated in fetal and adult human ACTH-stimulated cortex in Xing *et al.* 2010^38^. Expression = z-scored mean log2(TPM+1) of stimulated versus unstimulated mDZ. (**I**) GSEA of differential expression in stimulated versus unstimulated mDZ using KEGG Medicus terms and (**J**) GO biological process terms. (**K**) Heatmap highlighting distinctness of genes in GO Biological Process terms related to positive or negative cell cycle regulation. Heatmap colored by gene overlap (Jaccard similarity). Term order determined by hierarchical clustering, shown with dendrogram. (**L**) Z-scored expression averages for cell cycle related genes in stimulated and unstimulated mDZ. (**M**) Volcano plot of differential expression highlighting signaling pathway genes. (**N**) HES1 fluorescence intensity from IF in unstimulated versus stimulated day 32 mDZ. Dots represent images (n = 3 biological replicates). (**O**) Volcano plot highlighting inflammation-related terms present in differentially expressed gene list. All graphs in this figure show mean ± SEM. Abbreviations: AIC = ACTH, CRH, IGF2 stimulation; DZ = definitive zone; GSEA = Gene Set Enrichment Analysis; IF = immunofluorescence; mDZ = matured DZ; NES = Net Enrichment Score; TEM = transmission electron microscopy; TPM = transcripts per million.

### Transcriptional ACTH Response Recapitulates Normal Adrenal Function

To evaluate how stimulation impacts the transcriptional landscape of mDZ cultures we performed bulk RNAseq of d32 stimulated and unstimulated cells (**Fig. Supp. 5C**). Many transcripts related to steroidogenesis, cell signaling, and metabolism were significantly upregulated in the stimulated condition (*CYP21A2*, *HSD3B2*, *CYP11B1*, *MRAP*), while others were downregulated (*GADD45A*, *SCARA5*; **Fig. 3G, Fig. Supp. 5D, Table 1**). We also saw significantly upregulated expression of the steroidogenic enzymes *HSD3B2* and *CYP21A2* along with *MC2R* upon stimulation by RT-qPCR, while zone-specific gene expression was not impacted in either assay (**Fig. Supp. 5E, F**). Previous studies have characterized transcriptional responses of human fetal and adult adrenal cortexes to ACTH *in vitro*^38^. In our cultures, we tested expression of those transcripts that were found to be upregulated in both human adult and fetal adrenals upon exposure to ACTH and found that they are also upregulated here (z-scored expression averages, **Fig. 3H**; per-sample Z-scores, **Supp. Fig. 5D**). These similarities support that our adrenocortical model faithfully captures the human adrenal ACTH response. To characterize differential gene expression, we utilized Gene Set Enrichment Analysis (GSEA) to assess which known pathways and biological processes were most altered in the stimulated condition compared to unstimulated and vice versa. The KEGG Medicus database GSEA revealed terms reflective of robust ACTH response and indicative of a shift in metabolic state consistent with adrenocortical steroidogenesis, such as ACTH cortisol signaling pathway and cholesterol biosynthesis (**Fig. 3I**, **Table 2**). GSEA plots of running enrichment scores displayed steep leftward peaks demonstrating significant pathway activation (**Supp. Fig. 5G**).

Interestingly, top GO biological process GSEA results indicate an enrichment of terms related to both cellular division and cell cycle arrest (**Fig. 3J**, **Supp. Fig. 5H, Table 2**). This simultaneous increase and decrease in cycling related transcripts reflect the complex, dual role of ACTH. On one hand, ACTH drives terminal differentiation (which typically requires cell cycle arrest)^39,40^ and ACTH treatment of *in vitro* adrenocortical cultures have an anti-mitotic effect^41^. On the other hand, ACTH clearly acts as a mitogen *in vivo*, driving adrenal hyperplasia during chronic stress and in diseases such as congenital adrenal hyperplasia^36,41–43^. In health, this proliferative response may serve as a protective mechanism to maintain the progenitor pool when differentiation is highly active. However, while the structural development of the DZ is independent of ACTH, its functional maturation is not^39^. Thus, it remains unclear exactly which cell populations are proliferating upon ACTH exposure. We confirmed that the upregulated positive and negative cell cycle terms are transcriptionally distinct programs, rather than artifacts of redundancy between GO term gene lists **(Fig. 3K**). Because known proliferation markers did not meet significance threshold in the differential expression test, we examined Z-scored expression of cell cycle-related markers and observed positive z-scores for most proliferation markers (**Fig. 3L**). Beyond proliferation, we wanted to understand how stimulation alters cell lineage trajectories. To do this, we checked for the overrepresentation of zone-specific markers. We hypothesized that ACTH stimulation might enrich for a specific cellular identity. However, we observed that both DZ and FZ markers from the consensus DEG list were upregulated (**Supp. Fig. 5F**). This indicates that rather than cleanly shifting cells toward a purely FZ-like or DZ-like state, ACTH induces a more complex, overlapping transcriptional response.

To understand the mechanism by which stimulation alters cell identity, we examined the DEG list for changes in intercellular communication networks known to regulate adrenal zonation (**Fig. 3M**, **Table 1**). First, we observed a strong overrepresentation of the TGFb pathway (e.g., *BMPR2*, *LEFTY2*, *TGFBI*, *INHA*). This was an expected result, as the transition to stimulation media requires the removal of the TGF-β inhibitor SB. Next, we investigated pathways directly responsible for identity maintenance. We found that WNT4, which enforces zG/zF identity through mutual antagonism with PKA signaling^2,40,44^, was significantly upregulated in the stimulated condition. This aligns with physiological responses observed in primary tissue, suggesting WNT4 upregulation is a key mechanism for maintaining zonation in our model^45^. Conversely, core NOTCH signaling pathway members (*NOTCH1*, *JAG1*, HEY2) were downregulated. Because the NOTCH response gene HES1 did not meet the significance threshold, we assessed its expression at the protein level using IF. Using fluorescence intensity, we observed a modest decrease in HES1 expression in the stimulated condition, supporting that our finding of decreased NOTCH-related transcripts is indeed a reflection of overall decrease in NOTCH signaling (**Fig. 3N**). Finally, we observed a disproportionate number of inflammation-related transcripts that were significantly upregulated, indicating activation of gene regulatory programs that are implicated in mediating inflammation (**Fig. 3O**). Together, these data demonstrate that stimulation treatment impacts mDZ cells in two distinct ways. First, steroidogenesis transcripts are upregulated in one subset, potentially leading to terminal differentiation and suppression of mitotic programs. Second, it reshapes the signaling environment by suppressing NOTCH and promoting WNT4, pathways fundamentally implicated in the establishment and maintenance of definitive zone identity.

### Identification of Consensus Zonation Markers by Meta-Analysis of scRNAseq Expression Data

Many adrenal cortex and capsule markers are well understood to represent each cell type within the adrenal gland. Such ‘canonical’ markers are mostly derived from studies in animal models. However, they may not accurately represent the human fetal status, the time frame our cells fall into. For example, the mouse capsule marker *GLI1*^46–48^, has not been consistently observed to be a capsule marker in recent human studies^1,49^. Reports of capsule-specific GLI1 expression in humans are limited to 4-5 WPC, while at 8 WPC and later, GLI1 expression was identified in the cortex^7,50^, indicating a potential species-related difference. Furthermore, there are adrenal cell types whose gene expression profiles remain unclear, such as zG-resident progenitor cells. zF cells also have strong gene expression overlap with neighboring cell types, leaving few known markers with zF specificity. Together, these limitations highlight a need for clarity. Recently, a multitude of single cell transcriptomics analyses of human fetal and adult adrenal glands have been performed, offering an opportunity to establish a decisive marker signature for each adrenal cell type^1,7,28,29,49,51–53^. Thus, to define markers of cortex and capsule cell types, we performed a meta-analysis, combining published differential gene expression results from 9 publications (4 fetal, 5 adult; **Fig. 4A**, **Table 3**). We accounted for the relative nature of differential gene expression and the varying resolution of cluster annotations across the original datasets. Because neighboring adrenal cortex zones naturally share steroidogenic enzymes and signaling environments, we defined markers based on both regional specificity (aldosterone-secreting zG/DZ, progenitor zG/DZ, zF, and zR/FZ) and biological overlap (pan-cortex, pan-zG/DZ, shared zG+zF, shared zF+zR, shared capsule+zG). Pan-zG/DZ markers separate both aldosterone secreting cells and zG progenitors from the rest of the non-zG cortex, but do not distinguish between them. Similarly, pan-cortex markers separate the entire cortex from the capsule without zone-specific restriction. Shared capsule+zG markers instead highlight shared signaling niches between these adjacent, but highly distinct cell populations (**Fig. 4B**). While original authors did not always name an aldosterone synthase (*CYP11B2*) negative zG cluster “zG progenitor”, we assigned it this identity based on the known existence of zG progenitors^6,35,50^ plus the presence of the stemness markers *ID2*, *HES1* and *LEF1*^54–57^ and pan-zG/DZ markers, but absence of CYP11B2. Aldosterone-secreting cell markers included *CYP11B2*, *ETV1*, *COL15A1* (**Fig. 4C**, **Fig. Supp. 6**, red). Pan-zG/DZ markers included *DAB2*, *MAML3*, *GNAI1*, and *HOPX* (**Fig. 4C**, **Fig. Supp. 6**, orange). Shared zG+zF markers (*HSD3B2, ETV5*, *AKAP7, CCN3/NOV*; green) included those enriched in clusters identified by the original authors as in transition from zG towards zF as well as genes found in both zG/DZ and zF (but not in zR/FZ). Only 9 zF-specific genes (e.g., *DLK1*, *CTNNAL1*, *ABCC3*; teal) were identified. We also identified shared zF+zR markers (*CYP17A1*, *CYP11B1*, *SCARB1*; blue), zR/FZ-specific markers (*GSTA1*, *SULT2A1*, *CYB5A*; purple), and capsule markers (*C1R*, *MGP*, *IGFBP6*). Finally, we identified cell surface markers that warrant future investigation for their utility in sorting strategies, including *CD248* (capsule), *CD46* (pan-cortex), and *CD55* (zG+zF). The top 15 markers per cell type are shown in **Fig. 4C**; all markers are shown in **Fig. Supp. 6; Table 4**).

**Fig. 4:**
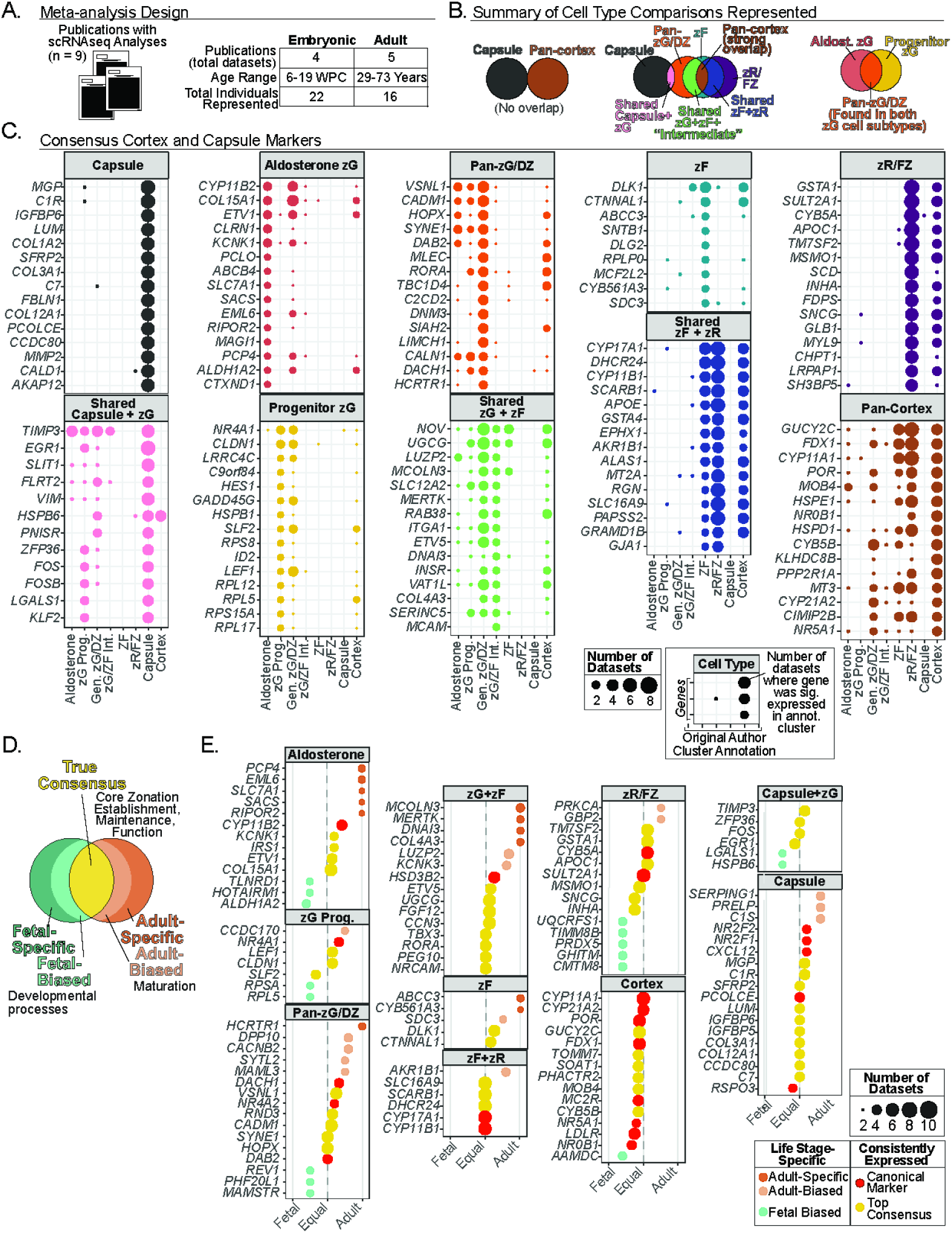
Identification of Human Adrenocortical Consensus Zonation and Maturation Markers. (**A**) Summary of meta-analysis study design: number of scRNAseq analyses included, number of individuals represented, and age ranges of individuals (**B**) Summary of cell types identified, comparisons between cell types, and expression overlaps characterized. Each expression overlap and distinct cell type is a “cell category”. (**C**) Dot plot showing the top 15 markers of each cell category. Cell category at the top of each dot plot corresponds to the comparison diagram in (**B**). Dot size = number of times each marker (y-axis) was present in a cell type cluster from the original publication (x-axis). (**D**) Diagram depicting interpretation of life stage specificity. (**E**) Diverging dot plot of adult or fetal expression bias. Established markers for each cell type (canonical; red); strongest conserved expression across life stages (yellow); fetal bias (teal; left of center); adult biased (orange; right of center) expression. Distance from center line indicates stronger bias. Size of dot represents number of times a gene was present in the cell type category the in the original datasets. Abbreviations: DZ = definitive zone; FZ = fetal zone; WPC = weeks post conception; zG = zona glomerulosa; zF = zona fasciculata; zR = zona reticularis.

**Fig. 5:**
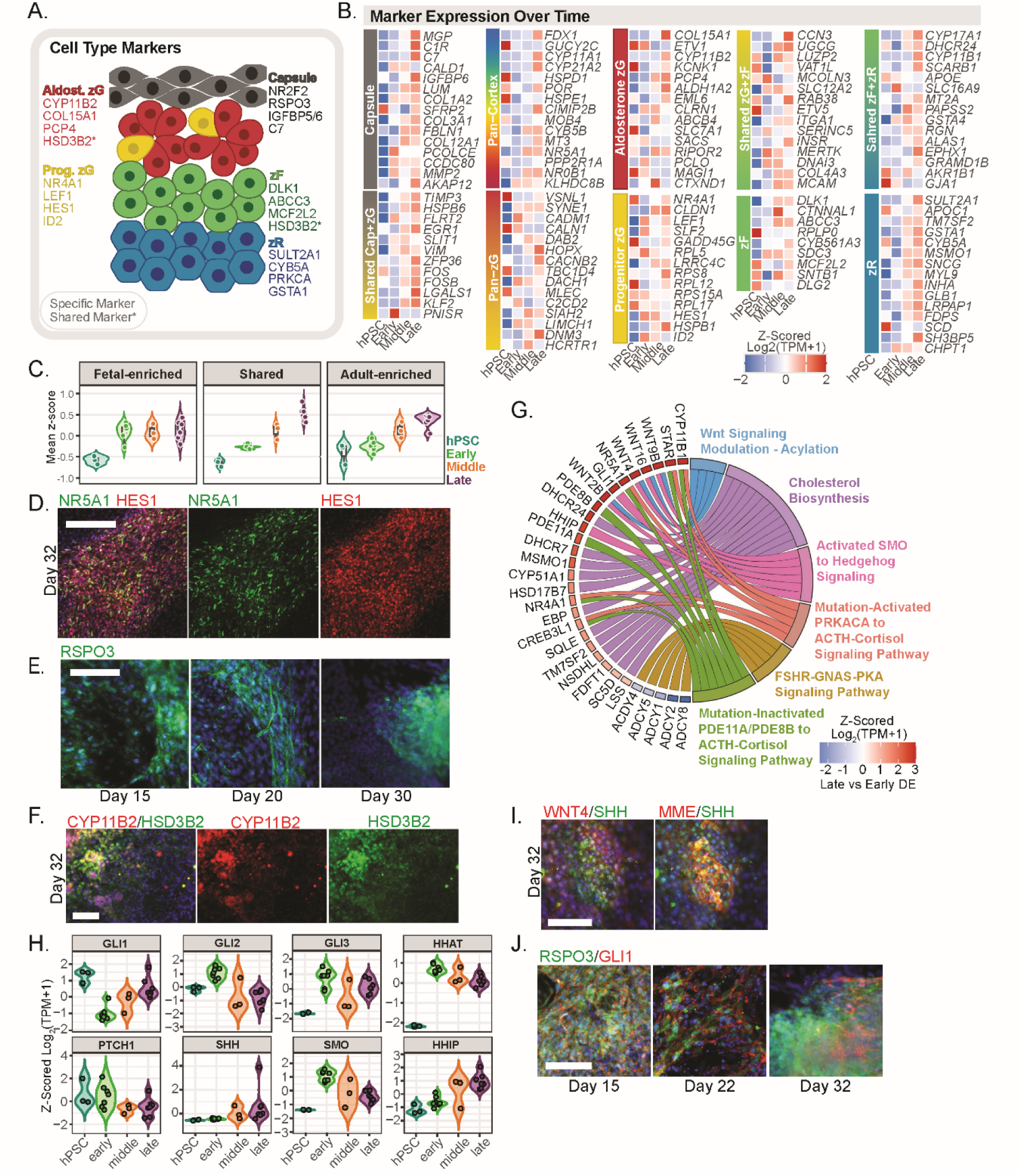
Cellular Subtype Characterization. (**A**) Diagram of adrenal zonation and consensus markers of distinct cell types. (**B**) Heatmap of z-scored expression (log_2_(TPM+1)) in mDZ of the top 15 markers for each cell type identified in Fig. 4B. (**C**) Averaged z-score for all fetal-enriched, shared, and adult-enriched markers identified in Fig. 4E. (**D**) IF for the cortex marker NR5A1 with the progenitor marker HES1. Scale = 200µm. (**E**) IF for the capsule marker RSPO3. Scale = 100µm. (**F**) IF for CYP11B2 and HSD3B2 alongside individual color channels. Scale = 100µm. (**G**) Chord diagram of KEGG Medicus terms enriched in GSEA analysis of late versus early mDZ differentiation. Genes enriched in each GSEA term with z-scored expression (log2(TPM+1), left), GSEA term (right). GSEA terms. (**H**) Violin plot of z-scored log2(TPM+1) expression in differentiating mDZ samples over time. Hedgehog pathway member expression is shown over time. (**I**) IF of unstimulated mDZ cells on differentiation day 30 for WNT4 and SHH, left; MME and SHH, right. (**J**) IF on days 15, 22, and 32 for RSPO3 and GLI1 in unstimulated mDZ cells. Scale =100µm. For all IF, DAPI was used as a nuclear counterstain. Abbreviations: DZ = definitive zone; FZ = fetal zone; GSEA = gene set enrichment analysis; IF = immunofluorescence; mDZ = matured DZ; TPM = transcripts per million; zG = zona glomerulosa; zF = zona fasciculata; zR = zona reticularis.

### Markers of Adrenal Maturation, Development, and Core Zonation Processes

Because our meta-analysis included both fetal and adult datasets, we evaluated the life-stage specificity of our identified markers. Cortex spatial organization is conserved from embryonic development throughout life with respect to which steroid each zone produces, and which signals are necessary to maintain this zonation^1,8,50,58,59^. We therefore expect that zone specific markers with equal representation across life stages are likely indispensable for zonation establishment, maintenance, and/or function. Conversely, markers with fetal specificity may have important roles in developmental processes, while adult-specific markers could represent maturation-associated processes (**Fig. 4D**). We identified adult-specific transcripts across multiple zones (*PCP4*, *SLC7A1*, in aldosterone secreting cells; *HCRTR1* in pan-zG; *ABCC3* and *CYB561A3* in zF). Adult-biased genes were present in almost all cell types, including the previously established maturation marker *AKR1B1* in shared zF+zR^60^. While no strictly fetal-specific markers were identified, several were fetal-biased (e.g., *ALDH1A2* and *TLNRD1* in aldosterone-secreting cells; *CMTM8*, *GHITM*, and *PRDX5* in zR/FZ), suggesting roles in zone-specific developmental pathways (**Fig 4E**, **Fig. Supp. 7A**, **Table 5**).

### Characterization of Represented Cell Types

The distinct expression profiles characterized in **Fig. 4** allowed us to characterize our own adrenocortical model system for the presence of these distinct cell types (**Fig. 5A**). Cells in the adrenal cortex are regenerated throughout life by the stem cell-like population residing in the capsule^6,46,61^, which differentiate into subcapsular progenitor cells that reside within rosettes alongside aldosterone-secreting cells^62,63^. These capsule cells are also critical for zonation and for secreting signaling cues that establish and maintain zonation, such as RSPO3^6,58^. We have shown that our cultures synthesize steroids of all three cortex zones (**Fig. 3A-C**). As capsule-derived signals are needed for aldosterone-secreting cell specification and identity maintenance^34,58^, we next sought to verify the presence of this capsular signaling program in our system. We analyzed transcriptional profiles throughout differentiation with bulk RNAseq across four differentiation stages (hPSC (day 0), early (days 8, 10), middle (day 15), and late (days 22-32; late samples were ACTH-stimulated) based on PCA clustering of differentiation over time (**Fig. 2C**). Z-scored expression of zonation markers identified in our meta-analysis (**Fig. 4B**) revealed that mDZ cultures successfully establish gene expression profiles corresponding to adrenal capsule and diverse adrenocortical cell types, including zG progenitors, aldosterone producing cells, zF, and zR (**Fig. 5B**). Next, we evaluated the transcriptional maturation signature. Adult-specific and-biased markers were grouped and defined “adult-enriched”. The strongest upregulation over time was observed in the shared category, containing genes that were expressed consistently throughout life, followed by adult-enriched genes. In both groups, average z-scores gradually slope up towards late differentiation. Fetal-enriched marker expression was not meaningfully upregulated, however. Overall, this suggests that our cultures transcriptionally resemble adult cortex and capsule more than fetal (**Fig. 5C**). We then validated these transcriptional profiles at the protein level by IF in unstimulated day 32 mDZ cultures. We confirmed the presence of these progenitors, showing overlapping expression of NR5A1 and HES1 (**Fig. 5D**), capsular cells expressing RSPO3 starting at day 15 (**Fig. 5E**), and aldosterone secreting cells co-expressing CYP11B2 and HSD3B2 (**Fig. 5F**). Together, these data demonstrate that mDZ cultures are heterogeneous, containing capsule cells, zG progenitors, and multiple distinct steroidogenic components.

### Analysis of Transcriptional Profiles and Zonation Pathways

Next, we sought to characterize the signaling networks driving mDZ culture differentiation. Differential expression analysis confirmed expected stage-specific transitions: early stages saw upregulation of EMT-related transcripts (*SNAI2*) and adrenal axial identity markers (*HOXA7*)^7^, while the late stages were characterized by upregulation of steroidogenic enzymes (*CYP11B1*; **Tables 5**, **6**). To identify core signaling networks active in mature cultures, we performed GSEA on the late versus early mDZ differential expression results (**Tables 5**, **6**). This revealed strong enrichment for terms related to ACTH-induced PKA signaling and cortisol production along with WNT and hedgehog signaling pathway activity, all of which are implicated in adrenal development^6,48,50^ and zonation maintenance^58,64^ (**Fig. 5G**, **Table 6**). We thus specifically investigated the roles of WNT and hedgehog signaling in our model. RNAseq confirmed Hedgehog and WNT signaling pathway member expression (**Fig. 5H**). By IF, we observed a subpopulation of WNT4+ cells co-expressing SHH. These SHH+/WNT4+ cells were also positive for the DZ marker MME, consistent with a more progenitor-like identity (**Fig. 5I**). Meanwhile, we also observed a population of RSPO3+/GLI1+ capsule cells across all timepoints (**Fig. 5J**). Together, these results indicate that hedgehog and WNT signaling between zG progenitors and capsule cells is functionally established in our *in vitro* system.

## Discussion

In this study we describe an *in vitro* model for human adrenocortical steroidogenic cells of all three adrenocortical zones that differentiate simultaneously alongside adrenal capsule cells from human pluripotent stem cells (**Fig. 1**). By adapting a protocol for gonadal directed differentiation, we first differentiated adrenocortical progenitor cells (AdCPs), followed by definitive zone (DZ) specification using a combination of TGF-b inhibition and B27 supplement, containing vitamin A (**Fig. 2**). This system allows for the differentiation of both aldosterone-secreting cells, cortisol-secreting cells, and androgen precursor secreting cells simultaneously by generating a coordinated, self-sustaining signaling niche comprised of WNT signaling (WNT4 and RSPO3), and reciprocal SHH signaling from DZ progenitors acting on capsular GLI (**Fig. 5H**-**J**). By generating all cortical zones, the subcapsular cortex cells, and the capsule cells in a single differentiation, our model is a fusion between organoid and 2D culture.

We found that TGF-b receptor inhibition is essential for DZ differentiation and that the timing of this inhibition is important, consistent with Mayama et al.^5^. Differentiating cells must establish AdCP identity prior to inhibiting TGF-b (**Fig. Supp. 2A**-**D**), but once past the progenitor stage, must continue to be inhibited throughout the course of differentiation (**Fig. Supp. 4 B**-**D**). In our system, the concurrent presence of capsular RSPO3 and cortical progenitor SHH suggests that zonation in this system arises through niche signaling interactions analogous to those observed *in vivo*. We also found that steroidogenic differentiation did not require extrinsic treatment with ACTH or other activation of cAMP/PKA (**Fig. Supp. 3C**, **D**), consistent with findings about ACTH’s role in cortex development^25^, and in agreement with the findings of Sakata *et a*l^25^ and Mayama *et al*^5^. Our mDZ cells demonstrated a robust functional response to ACTH, making them highly suitable for disease modeling. Stimulation induced 16-fold increase in cortisol secretion (**Fig. 3D**) as well as upregulation of combined with a coordinated enzymatic shift (upregulation of *CYP21A2* and *HSD3B2*, key cortisol synthesis enzymes; downregulation of *HSD11B2*, which converts cortisol to an inactive form) leading to an increase in biologically available cortisol. Furthermore, stimulation induced characteristic transcriptional changes of ACTH response in human primary cortex cultures^38^ (**Fig. 3F**, **Fig. Supp. 5D**). Together, these transcriptional changes paint a picture of robust functional response to ACTH.

Our meta-analysis of capsule and cortex cell markers established a comprehensive roadmap of gene expression signatures for adrenal cortex cell types, bridging fetal and adult profiles (**Fig. 4C, E**). These gene expression signatures strengthened the validation of our differentiation strategy, while also providing a resource that can be utilized for the identification and prospective isolation of specific adrenal cortex cell populations, the benchmarking of future differentiation protocols, and the study of adrenal cortex renewal. We identified consensus markers both for established canonical populations and for previously poorly defined populations, most notably the aldosterone-secreting zG and zG progenitor cells. zG progenitors act as cell reservoirs, serving as a source of adrenal renewal^50^. Understanding what maintains these progenitor cells in an undifferentiated state, and how they exit this state, are invaluable for understanding how the adrenal cortex sustains itself under chronic demand, and for understanding disruptions of self-renewal in aging and cancer^65^. Here, we identified zG progenitor population markers that are downstream transcriptional mediators of signaling pathways such has *HES1* (NOTCH), *LEF1* (WNT), and *ID2* (BMP/TGF-b) (**Fig. 4C**), all of which are strongly associated with stemness in a variety of cellular contexts^54–57^.

Collectively, our data and existing literature implicate the NOTCH1/JAG1 axis as a primary gatekeeper of the adrenocortical progenitor pool, maintaining these cells in an undifferentiated state. Under baseline conditions, zG progenitors co-express downstream NOTCH effectors (*HES1*, *MAML3*; **Fig. 4C**, **Table 4**). However, when stimulated to differentiate with ACTH, CRH, and IGF2, key upstream NOTCH members (*JAG1*, *NOTCH1*, *HEY2*) are significantly downregulated (**Fig. 3M**, **Table 1**). This aligns with physiological development, where repressing NOTCH signaling prevents DZ cell differentiation^5^ and is associated with steroidogenic differentiation^5,25^, while NOTCH hyperactivity is linked to poor cortical organization and adrenocortical carcinomas^65–67^. Furthermore, the AdCP to FZ developmental transition has been shown to include downregulation of *HES1*, indicating that it poises progenitor cells for steroidogenic differentiation^7^. The regulation of this stemness network is complex. We found that *HES1* was not meaningfully downregulated upon stimulation, despite the overall negative shift in NOTCH signaling (**Fig. 3M**, **N**; **Table. 1**). HES1 may instead be sustained by other mechanisms, such as LEF1^56^, also identified in this study as zG progenitor-specific. Unexpectedly, our meta-analysis revealed that the non-canonical (indirectly-acting^68^) NOTCH signaling modulator DLK1, well established as essential for adrenocortical development^7^, was identified as zF-specific. Meanwhile, a NOTCH1 signaling activator, *CCN3* (encoding NOV), was common to both zG and zF. DLK1 has previously been reported as a marker of a subcapsular, relatively undifferentiated population that becomes more abundant with age and is mutually exclusive with aldosterone-secreting cell clusters^69^. Additionally, DLK1 mutations are frequently observed in adrenocortical carcinoma, serving as a predictor of malignancy, but are not observed in aldosterone-producing adenomas^70^. Finally, DLK1 overexpression in H295R has been shown to upregulate markers of shared zF+zR and zR while downregulating transcripts identified in this study as specific to aldosterone-secreting cells^70^. Worth noting in this context is the fact that cortisol levels also rise with age, while hedgehog and WNT signaling decrease^52^, as do DHEA-S and aldosterone^18^. In light of this information, we propose that DLK1 may mark a distinct cortical zF progenitor population in the human adrenal. Altogether, these findings highlight the importance of NOTCH signaling in adrenocortical progenitor maintenance.

In all, this mDZ differentiation strategy establishes a powerful new paradigm for studying the human adrenal cortex and capsule. Our *in vitro* system generates the niche responsible for the simultaneous development of all three cortical zones and the adrenal capsule in a single culture, providing a scalable and biologically accurate platform for investigating mechanisms of human adrenal development, renewal, aging, and oncogenesis. Furthermore, the identification of distinct progenitor reservoirs and their governing molecular regulators, specifically within the NOTCH and WNT signaling axes, offers critical new insights into adrenal development and zonation.

## Limitations of the Study

While we characterized the transcriptional landscape of differentiating adrenocortical cells *in vitro* through bulk RNAseq, RNA sequencing at the single cell level in the future will be informative towards understanding finer details of differentiation mechanisms. Additionally, the heterogeneous nature of the culture, while reflective of developmental dynamics, complicates assignment of cell-intrinsic versus niche-mediated effects. Furthermore, it will be important to determine whether our mDZ cultures have islands of distinct cell types or whether they are intermixed.

## Table Legends

**Table 1:** Differential expression data of stimulated versus unstimulated mDZ. Abbreviations: mDZ = matured DZ.

**Table 2:** GSEA results for stimulated versus unstimulated mDZ. Abbreviations: GSEA = gene set enrichment analysis.

**Table 3:** Original manuscript and donor data for meta-analysis. Includes DOIs of original publications and number of individuals represented as well as sexes and age ranges for individuals in publication.

**Table 4:** Data corresponding to consensus zonation marker and temporal bias identified in meta-analysis.

**Table 5:** Differentially expressed genes between stages of differentiation into mDZ from hPSC (day 0) to late (days 22 and 32). Includes differential expression between the following stages: hPSC versus early; early versus middle; late versus middle; late versus early. Abbreviations: hPSC = human pluripotent stem cell; DZ = definitive zone; mDZ = matured DZ.

**Table 6:** Late versus early mDZ differentiation GSEA results. Abbreviations: GSEA = gene set enrichment analysis.

**Table 7:** Antibody product information and primer sequences for all antibodies and primers used in this study.

## Resource availability

### Lead contact

Further information and requests for resources and reagents should be directed to Nadja Zeltner.

### Materials availability

Research materials generated in this article will be distributed upon reasonable request and upon completion of Material Transfer Agreements (MTAs).

### Data and code availability

All data generated or analyzed in this study are included in this article and its supplementary data file. Raw data points are available from the corresponding author on a reasonable request. RNA-seq data will be accessible to reviewers on GEO under accession GSE331200 with a reviewer token. Upon publication, the data will be made available to the public. This paper does not report original code. Raw data will be deposited on Figshare or a similar resource before publication.

### Author Contributions

N.Z. and J.M. conceived and designed the experiments; J.M., C.J., B.D. conducted experiments; J.M., C.J., K.T, T.K., and N.Z. analyzed and interpreted the data; J.M. and N.Z. wrote the manuscript; N.Z. provided mentoring and financial and administrative support and approved the final version of this manuscript.

## Acknowledgements

We wish to thank Tripti Saini for her critical reading on this manuscript and Dr. Ya-Wen Chen for her mentorship. The authors gratefully acknowledge use of the services of the Georgia Electron Microscopy core facility, 302 E. Campus Rd, Athens, GA 30602. This research was supported by NIH/NICHD to Zeltner 1R01HD115812-01A1.

## Experimental model and study participant details

### Human Cell lines

Experiments in this study were carried out using the human embryonic stem cell (hESC) line MEL1 (hPSC-ctr-MEL1, NIH registry #0139).

## Methods

### Human Pluripotent Stem Cell Culture

Human pluripotent stem cells (hPSCs) were maintained as described previously^71^. Briefly, hPSCs were grown in colonies on rh-Vitronectin (VTN; Invitrogen A31804) in Essential 8 Media (E8; Gibco A1517001) and passaged every 3-4 days by dissociation with EDTA.

### Adrenocortical Cell Differentiation

Adrenocortical progenitors (AdCPs) were differentiated by dissociating hPSCs for 10min in EDTA solution into single cells and subsequently plating them in E8 media with ROCK inhibitor (Y-27632, 10µM; Biogems 1293823) at a density of 10,000 cells per cm^2^ in 24-well TC-treated plates (Corning 3527) coated with VTN at 5 μg/mL for 1 hour at room temperature. After 24 hours, on day 0, E8 was replaced with 0.5mL per well Essential 6 media (E6; Gibco A1516401) containing 4µM Chir99021(Chir; Tocris 4423). Chir media was replaced each day twice more for a total of 3 days of Chir media treatment. Starting on day 3, media was replaced daily for a total of 3 days with 1.0mL per well E6 media containing 20ng FGF9 (R&D Systems 273-F9-MTO) and heparin (1:2000, StemCell Technologies 7980). Starting on day 6, media was replaced every other day with 1mL per well E6 media. On day 10, media was changed to 1.0mL per well of E6 with 10µM SB431542 (Tocris 1614) and 1:50 B27 supplement with vitamin A (Gibco 17504-044) and fed every other day.

### Gonad and Kidney Cell Differentiations

All protocols were performed according to original publications^26,31^ by seeding cells as described above and performing daily media changes unless otherwise specified. Briefly, for gonad differentiation, cells were incubated in E6 media containing 3µM Chir for 4 days followed by a 3-day treatment with E6 containing 10ng/mL BMP4 (R&D Systems 314-BP) and 100ng/mL FGF9. For kidney differentiation, Chir concentration was increased to 6µM and no BMP4 was added.

### Steroidogenic Stimulation

On days 15, 20, or 30 of differentiation, media was replaced with 0.5mL of stimulation media comprised of E6 media containing 10nM ACTH 1-39 (ACTH; Tocris 3492), 40pg/mL CRH (R&D Systems 1151), and 25ng/mL IGF2 (R&D systems 292-G2) for 48 hours. ACTH and CRH concentrations were determined based on basal plasma ACTH and CRH during early development. ACTH 1-39 was selected as the stimulus for cortisol release because it represents a more physiologically relevant stimulation method than small molecules such as 8-br-cAMP or forskolin.

### RNA Extraction and RT-qPCR

RNA samples were collected with Trizol (Invitrogen 15596018) and purified with phenol-chloroform extraction. Isolated RNA was resuspended in nuclease-free ultrapure H2O (Invitrogen 10977015). cDNA synthesis was performed using 1 μg of RNA (iScript, Bio-Rad 1708841) on an Eppendorf X50s thermal cycler. RT-qPCR was performed using SYBR Green Supermix (Bio-Rad 1725272) with a Bio-Rad CFX96 Real-Time System and C1000 Touch thermal cycler and analyzed by CFX Maestro 1.1 (Version 4.1). Calculations were performed as relative fold change (2^-ΔΔCt^) with respect to MEL1 hESCs and expression of the housekeeping gene GAPDH. For primers used in this study please refer to **Supp. Data 7**.

### Immunofluorescence

Adherent cells were washed once in DPBS (PBS; Corning 21-031-CM), fixed in cold 4% paraformaldehyde (PFA, Thermo Scientific AAJ19943K2) for 10 minutes, and washed 3 times in PBS. Cells were then permeabilized in a solution of 0.3% Triton X-100 (Sigma X100) for 5 minutes and subsequently blocked in solution containing 0.2% Tween 20, 3% donkey serum (Sigma S30), 1% BSA (Sigma A4503), 0.01% sodium azide (Sigma S2002) in PBS for 40 minutes. Primary antibodies were diluted in antibody buffer (same as blocking buffer without tween-20) and cells were incubated overnight at 4°C. On the following day, cells were washed with PBS 3 times. Secondary antibody solution was prepared by diluting secondary antibodies at 1:400 and DAPI at 1:1000 into antibody buffer. Cells were incubated in secondary antibody solution while protected from light for 1 hour at room temperature and then washed 3 times in PBS. For antibodies used in this study please refer to **Table 7**. Fluorescence microscopy was performed on an Agilent Lionheart microscope. Image acquisition was performed using Gen5 software. Image quantification was performed on raw images in FIJI. Color overlays were generated in Adobe Photoshop.

### ELISA

ELISAs were performed for cortisol (Arbor Assays K003-H1), aldosterone (Arbor Assays K052-H1), and DHEA-S (Arbor Assays K054-H) following the manufacturer instructions. All media samples were frozen at-80C immediately after collection. ELISA kits with a 24-or 48-hour protocol option were all performed using the 48-hour option. ELISAs were read and analyzed on a BioTek Synergy 2. Calculations were performed by calculating %B/B0 in Excel and interpolating concentration using 4 parameter logistic regression (4PL) and nonlinear fit to the standard curve in GraphPad Prism.

### RNA sequencing

RNA was isolated as described above. RNA was subject to quality control by RIN measurement and samples with RIN < 6 and a minimum of 1×10^7^ reads were selected. mRNA transcripts with poly(A) tails were selected, cDNA synthesis was performed and library preparation was performed with NEBNext Ultra II Directional kit. Sequencing was performed with a depth of 40 million paired end reads using an Illumina HiSeq 2×150 at Admera Health.

### Meta-Analysis

Meta-analysis was performed in R v4.4.0^72^. Consensus gene lists were constructed from publicly available differential expression data from human adrenal scRNAseq experiments (**Table 3**). Gene symbols/aliases were first converted to their respective Ensembl identifiers and then “canonicalized” by conversion back to gene symbol using org.Hs.eg.db and AnnotationDbi^73^. Each time a gene appeared as differentially expressed in a dataset, its presence was considered a “vote”.

#### Consensus Marker Classification

##### Original Dataset Uniformization

To identify highly specific markers of adrenal cortex and capsule cell types, we developed a hierarchical, rule-based classification algorithm. Genes were first assigned to discrete categories based on their original cluster annotation: capsule, zona glomerulosa (zG), intermediate (zG to zF transition cluster), zona fasciculata (zF), zona reticularis (zR). In cases where there were multiple zG clusters, they were further subdivided into aldosterone secreting (based on the expression of *CYP11B2*) or zG progenitor (based on the presence of known zG markers but absence of *CYP11B2*).

##### Cell Type Category Construction

A stringent thresholding strategy was applied to ensure specificity. For example, zG progenitor and aldosterone secreting cells each required high detection scores (>= 2 to 3 votes) coupled with negligible signals in the capsule or other cortical layers (<= 1 vote). The exact R script containing the cell type category classification is as follows:

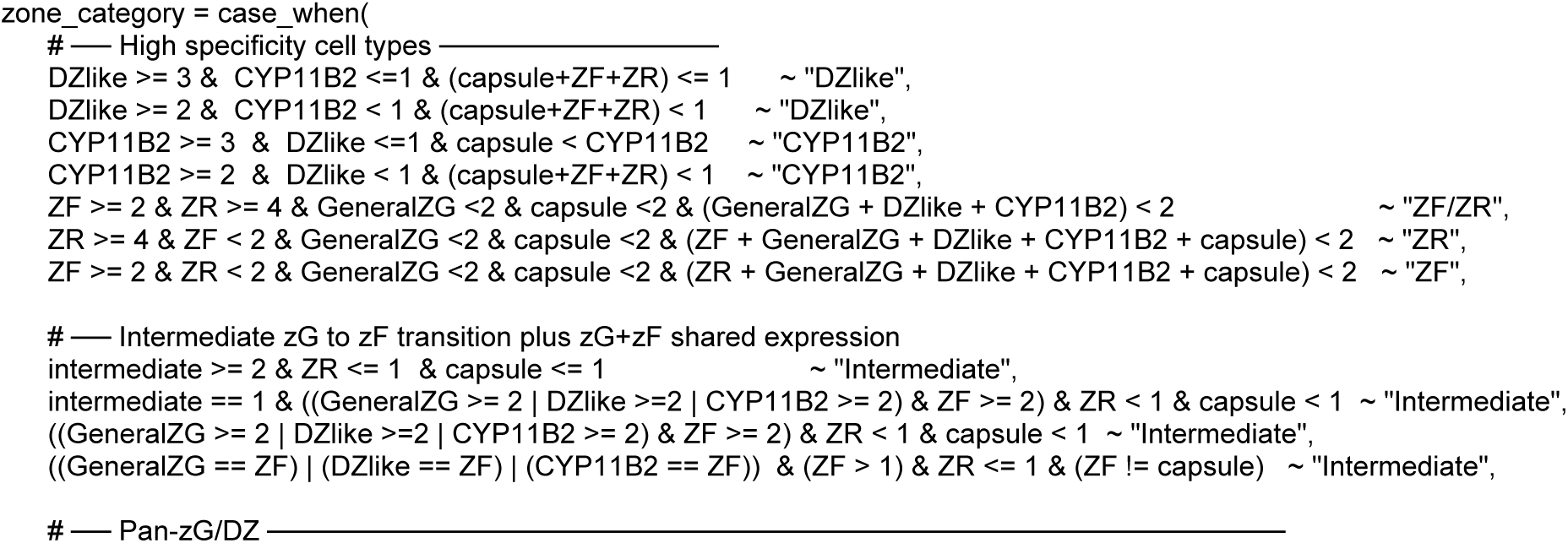
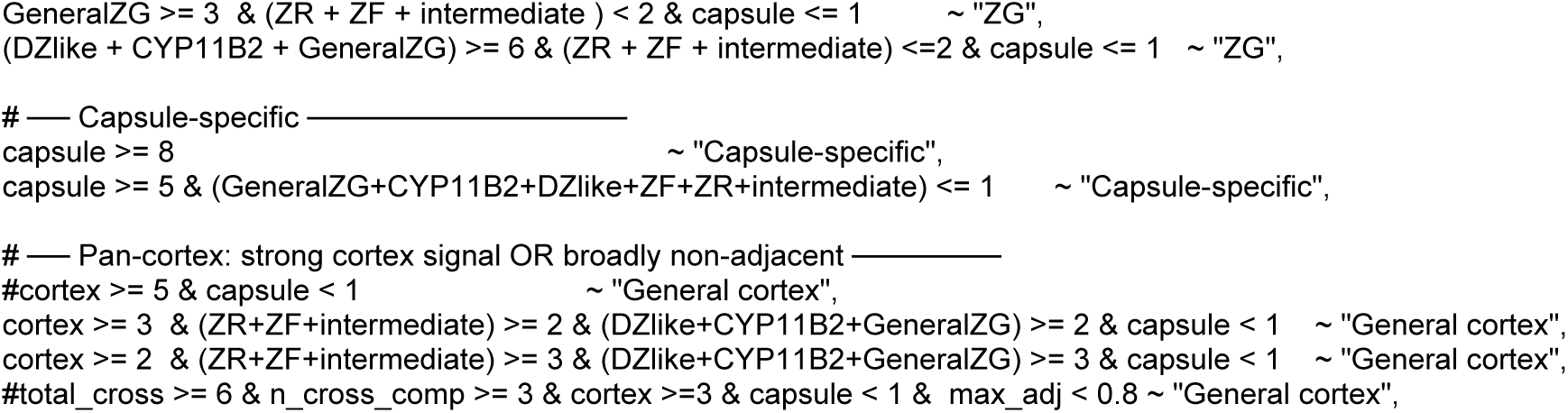

Genes failing to meet any of these mutually exclusive criteria were unclassified and removed from downstream marker analysis.

##### Marker Prioritization and Matrix Generation

Following zonal categorization, each gene was assigned a “defining vote” score corresponding to the primary zone to which it was classified. Consensus markers were ranked by the number of datasets it appeared in an annotated cell type but not those considered to be mutually exclusive. As a tiebreaker, we then sorted by average log2 fold change for each gene across all datasets, except for the Descartes adrenal cortex and capsule datasets because log2FC values were not available.

##### Temporal Specificity Calculation

Ages of fetal datasets range from 6-19 WPC, while adult datasets range from 29-73 years. In total, 22 embryos and 16 adults are represented in this analysis (**Table 3**). For life stage bias evaluation, marker genes were first classified by their stage specificity using a multi-dataset cross-reference approach: genes detected as zone-enriched in two or more adult adrenal datasets (n=5) but present in 0 fetal datasets were designated “adult-specific”. Genes present in greater than 50% of adult datasets but fewer than 25% of fetal datasets were called “adult-biased”. Genes enriched in in greater than 50% of fetal datasets but fewer than 25% of adult datasets were designated “fetal-biased”. Genes present in the consensus gene list but without meeting criteria described above were called “Shared”.

### RNA sequencing analysis

The following were all performed in R 4.4.0^72^ with dplyr^74^.

### Processing

RNAseq data was quantified by quasi-mapping using Salmon^75^ and annotated using txdbmaker^76^ and AnnotationDbi^73^ with GENCODE GRCh38 reference. Both raw counts and transcripts per million (tpm) were exported for downstream analysis.

### Z-scored Expression

Transcript expression was evaluated from tpm values. First, tpms were log-normalized by log_2_(tpm + 1). Z-scores were then calculated with the R function scale: t(scale(t(log_tpm), center = T, scale = T)).

### Differential Expression

Differential expression testing was performed with the R package DESeq2^77^. Raw counts were tested using the DESeq function. Test type used was likelihood ratio test (LRT) and fitType = local. Raw count matrices were modeled using a generalized linear model with a negative binomial distribution, incorporating differentiation batch as a covariate to control for differentiation-to-differentiation variability.

### Principle Component Analysis

For visualization and exploratory analysis, variance-stabilized expression values were obtained using the variance stabilizing transformation (VST) implemented in DESeq2. Principal component analysis (PCA) was performed on the transformed data to assess global transcriptional structure and sample relationships.

### GSEA

Gene Set Enrichment Analysis (GSEA) was performed using the GSEA function in ClusterProfiler^78^ using ENSEMBL gene IDs. Databases were prepared using msigdbr^79^. Unthresholded gene list was ranked by multiplying the sign of log_2_ fold change by - (log(pvalue)).

### Cell Cycle Pathway Distinctness

Overlap of genes driving GSEA term enrichment scores was determined with Jaccard matrix score calculation. Redundancy among enriched mitotic gene sets was assessed by calculating pairwise Jaccard similarity coefficients (intersection over union of core enrichment gene lists). This pairwise computation was executed across all terms to generate a symmetric similarity matrix tracking gene overlap. Pathways were organized with hierarchical clustering using Ward’s minimum variance method (ward.D2). Rows and columns were annotated by functional category (positive vs. negative cell cycle regulation) and the normalized enrichment scores (NES) calculated previously by GSEA.

### Assessment of Maturation State

To assess the maturation state of differentiating cells, we derived gene signature scores from markers identified as either biased towards embryo or adult, or consistent across all life stages (**Fig. 4E**, **Fig. Supp. 7**, **Table 4**). Genes detected as zone-enriched in two or more adult adrenal datasets (n=5) but absent from embryonic datasets were designated “Adult-enriched,” while genes enriched in two or more embryonic datasets (n=5) but absent from adult datasets were designated “Embryonic-enriched.” Genes with supporting evidence from both adult and embryonic datasets were classified as “Shared.” Genes appearing closest to the center and with the most votes for that cell category. For each bias class, a signature score was computed per sample as the mean z-scored log₂(TPM+1) expression across all member genes. Z-scoring was performed per gene across all samples prior to averaging, such that each gene contributed equally regardless of its absolute expression level. Signature scores were then compared across differentiation stages (hPSC, early, middle, late) to evaluate whether cells progressively acquired adult-type adrenal identity over the course of differentiation.

## Data Visualization/Plotting

RT-qPCR data and ELISA data were plotted using GraphPad Prism v11.0.0. The following R packages were utilized as described: Volcano plots were generated with EnhancedVolcano^80^. Heatmaps and dotplots (except for GSEA dotplots and Jaccard heatmap) were generated with ggplot2^81^. GSEA running enrichment plots were generated with GseaVis^82^. GSEA dotplots were generated by ClusterProfiler^78^. Chord diagram was constructed from differential expression results and KEGG medicus terms using SRplot^83^. Jaccard overlap heatmap for defining distinctness of gene enrichment in cell cycle GSEA was plotted with pheatmap^84^. Color schemes were generated with wesanderson^85^, viridis^86^, RColorBrewer^87^, and hclwizard.

### Transmission Electron Microscopy

Cells were prepared for transmission electron microscopy (TEM) and imaged as described previously. Images were acquired with accelerating voltage 200KeV^88^. TEM was performed by the Georgia Electron Microscopy core facility at UGA.

### Flow Cytometry

Cells were analyzed for MME expression by FACS as follows:

### Dissociation

Day 32 DZ cells were dissociated with 1 wash of PBS followed by a 30-minute treatment with Accutase (Innovative Cell Technologies AT104500) at 37°C. After 30 minutes, trypsin-EDTA solution (Sigma # T4174) was added at approximately 1:5 and incubated in 5-minute increments in 37C without shaking until individual cells could be observed detaching from one another under the microscope. Once cells were detached, they were collected in FACS buffer comprised of 10% fetal bovine serum (Corning 35-010-CV) in DMEM media (Gibco 11965118), quantified, and centrifuged at 4°C at 300 rcf for 3 minutes.

### Immunolabeling

After removing the supernatant and straining through a 40µm strainer (Falcon 352235), a minimum of 0.5×10^6 cells were resuspended in FACS buffer at 0.5&10^6 cells per 100uL. Samples containing both the primary and secondary antibody were treated with 5 µg/mL CD10 antibody (eBioscience 14-0106-82) for 30 minutes on ice. Primary-stained and secondary-only controls were incubated with 4 µg/mL goat anti-mouse IgG2b 488 (Invitrogen #A21141) for 30 minutes on ice. All antibody treatments were followed with two washes by centrifugation as described above. Unstained control samples were not treated with any antibody.

### Flow Cytometry

Cells were analyzed with a CytoFLEX (Beckman Coulter). Gating was performed by first excluding the signal from unstained control cells and then by excluding secondary-only control cells. Data were analyzed with FlowJo V10.10.1.

Diagrams were created with Biorender and Adobe Illustrator.

