## Supplemental figures for "Organoid-like Functional Adrenal Gland Cortex Derived From Human Pluripotent Stem Cells"

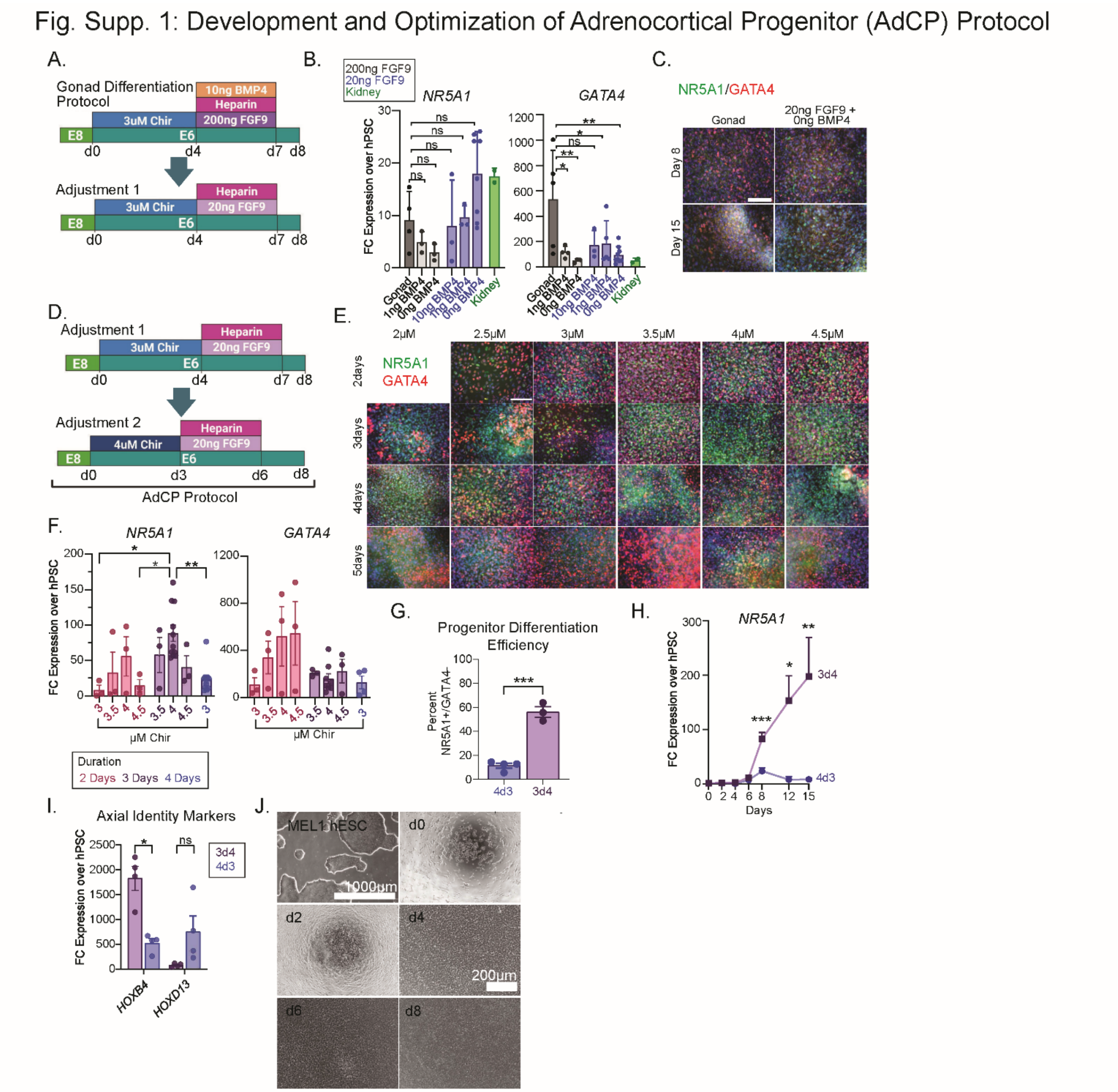


**Fig. Supp. 1: Development and Optimization of Adrenocortical Progenitor (AdCP) Protocol**
(**A**) Diagram of changes made to BMP4 and FGF9 concentrations from bipotential gonad differentiation described in Knarston *et al* 2019[26] protocol. (**B**) RT-qPCR of *NR5A1* and *GATA4* expression at day 8 using varying concentrations of BMP4 at 200ng FGF9 and at 20ng FGF9. Fold change = $2^{-\Delta\Delta Ct}$ relative to MEL1 hESC at day 0. One-way ANOVA with Dunnet's multiple comparisons (n = 3-5; *GATA4*, *p = 0.0159, all others n.s.). (**C**) Expression of the AdCP marker NR5A1 and gonad progenitor marker GATA4 at days 8 and 15 in the gonad differentiation protocol and with 20ng FGF9 + 0ng BMP4 detected by IF with nuclear counterstain DAPI (blue). (**D**) Diagram of changes made to Chir concentration and duration. (**E**) IF and (**F**) RT-qPCR of NR5A1 and GATA4 at each Chir concentration and duration tested. One-way ANOVA (*NR5A1*: 2d3 versus 3d4, *p = 0.016; 2d4.5 versus 3d4, *p = 0.038; 3d4 versus 4d3, **p = 0.001; all others n.s. *GATA4*: n.s.). (**G**) Quantification of AdCP differentiation efficiency by IF quantification, measured as percent of DAPI positive nuclei also positive for NR5A1, but not GATA4. Unpaired t-test (p*** = 0.0001). (**H**) RT-qPCR for *NR5A1* at 3 days of 4µM (3d4µM) Chir and 4 days of 3µM (4d3µM) Chir over time. Welch's t-test. (d8, ***p = 0.0004, n = 4-12; d12, *p = 0.028, n = 3-6; d15, **p = 0.0074, n = 6-9). Values for 3d4 and 4d3 conditions at day 8 are the same as those shown in (**F**). (**I**) Expression of adrenocortical HOX gene *HOXB4* and gonad HOX gene *HOXD13* at day 8 in 3d4µM Chir and 4d3µM Chir conditions. Multiple unpaired t-tests with Holm-Šídák correction (*HOXB4*, *p = 0.0045). (**J**) Phase contrast images of AdCP differentiation over time. All graphs in this figure show mean ± SEM. Abbreviations: AdCP = adrenocortical progenitor; Chir = Chir99021; hESC = human embryonic stem cell; IF = immunofluorescence.

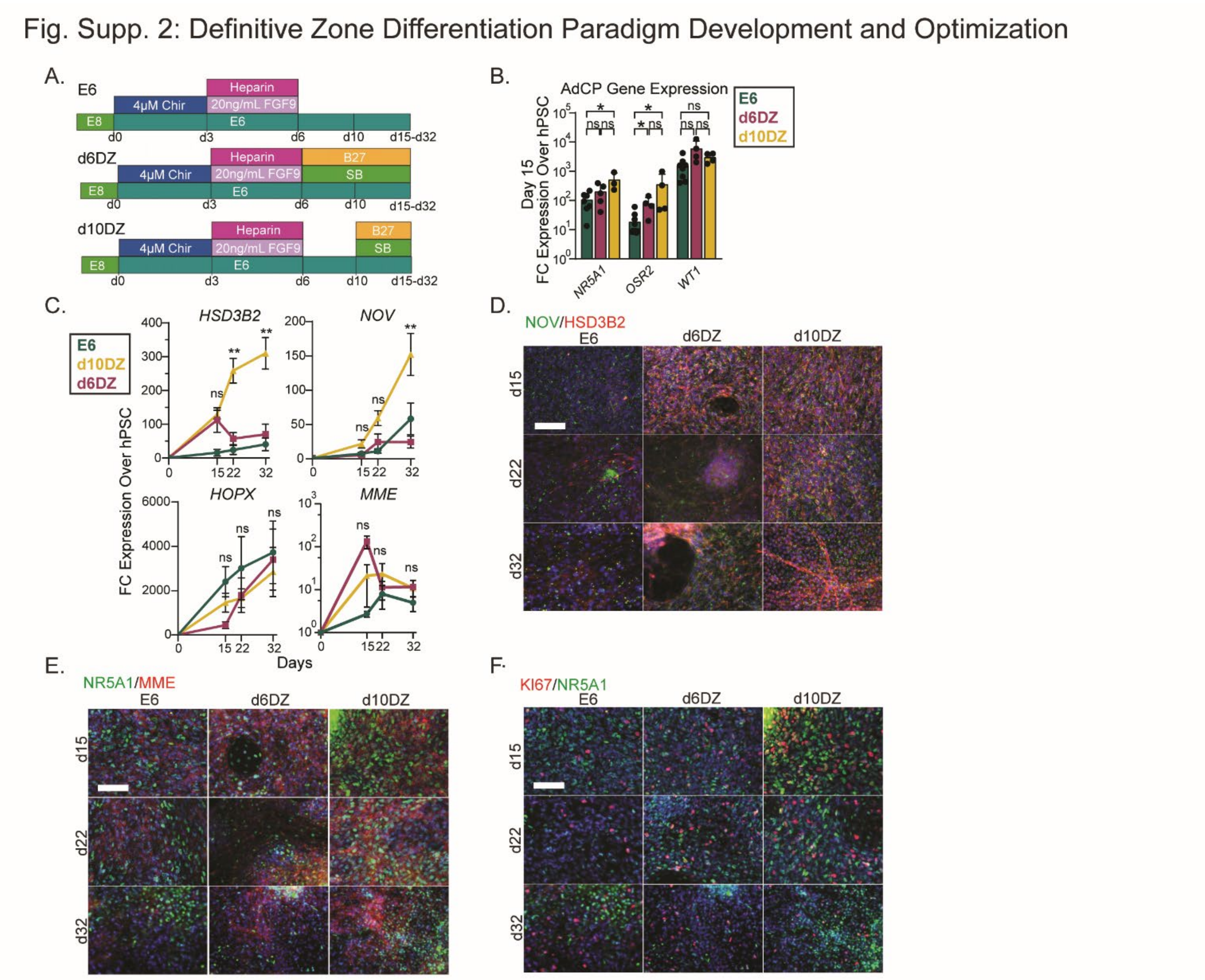


**Fig. Supp. 2: Definitive Zone Differentiation Paradigm Development and Optimization**

(**A**) Diagram of differentiation protocols tested for DZ differentiation paradigm generation. Cells were treated either with only E6 from day 6 or with SB and B27 starting at day 6 (d6DZ) or day 10 (d10DZ). (**B**) Expression of key AdCP markers at days 8 and 15 (RT-qPCR; *NR5A1*, n= 4-10; *OSR2*, n = 3-16). Unpaired t-test and Holm-Šídák multiple corrections (*NR5A1*: E6 versus d10DZ, *p = 0.046; *OSR2*: E6 versus d10DZ, *p = 0.048; E6 versus d6DZ *p = 0.016). (**C**) RT-qPCR for DZ markers over time (*HOPX*, n = 4-8; *HSD3B2*, n = 3-6; *MME*, n = 3-6). One-way ANOVA with Tukey's multiple comparison's test. Significance shown on graph corresponds to d10DZ vs d6DZ. Significant results of other comparisons between d10DZ and E6 or between d6DZ and E6 included here: (*HSD3B2* day 22: d6DZ vs. d10DZ ***p adj. = 0.0006; E6 vs. d10DZ p adj. = 0.0003; *HSD3B2* day 32: **d6DZ vs. d10DZ p adj. = 0.0013; E6 vs. d10 DZ p adj. = 0.0008; *NOV*, day 32: d10DZ vs d6DZ, *p adj. = 0.014; all other comparisons n.s.). (**D**) Expression of DZ markers (NOV, HSD3B2, MME) and (**E**) co-localization of MME with adrenocortical marker NR5A1 at days 15, 22, and 32 of differentiation by IF. Scale = 100μm. (**F**) IF for the proliferation marker KI67 and colocalization with NR5A1 at days 15, 22, and 32 in E6 (left), d6DZ (middle) and d10DZ (right). Scale = 100μm. All graphs in this figure show mean ± SEM. Abbreviations: AdCP = adrenocortical progenitor; B27 = B-27 supplement; DZ = definitive zone; E6 = base media; IF = immunofluorescence; SB = SB431542.

Fig. Supp. 3: Definitive Zone Differentiation Characterization Extended Data

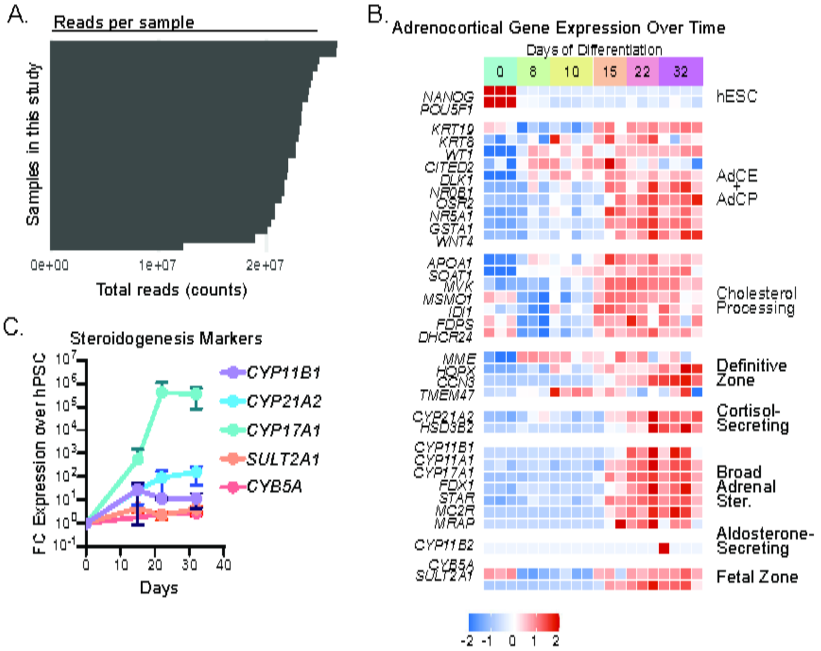


**Fig. Supp. 3: Definitive Zone Differentiation Characterization Extended Data**
(**A**) RNAseq quality control. Total reads per sample for all samples in this study (day 0, n = 3; day 8, n = 4; day 10, n = 3; day 15, n = 3; day 22, unstimulated, n = 3; day 22, stimulated, n = 3; day 32, unstimulated, n = 4; day 32, stimulated, n = 4). (**B**) Relative expression (Z-scored $\log_2$(TPM+1)) of development and zonation markers from **Fig. 2D** in individual RNAseq samples. Colors correspond to timepoint. (**C**) Steroidogenic potential over time was characterized in the DZ cultures by RT-qPCR for steroidogenesis mediating enzymes (n = 2-6). Abbreviations: DZ = definitive zone; TPM = transcripts per million.

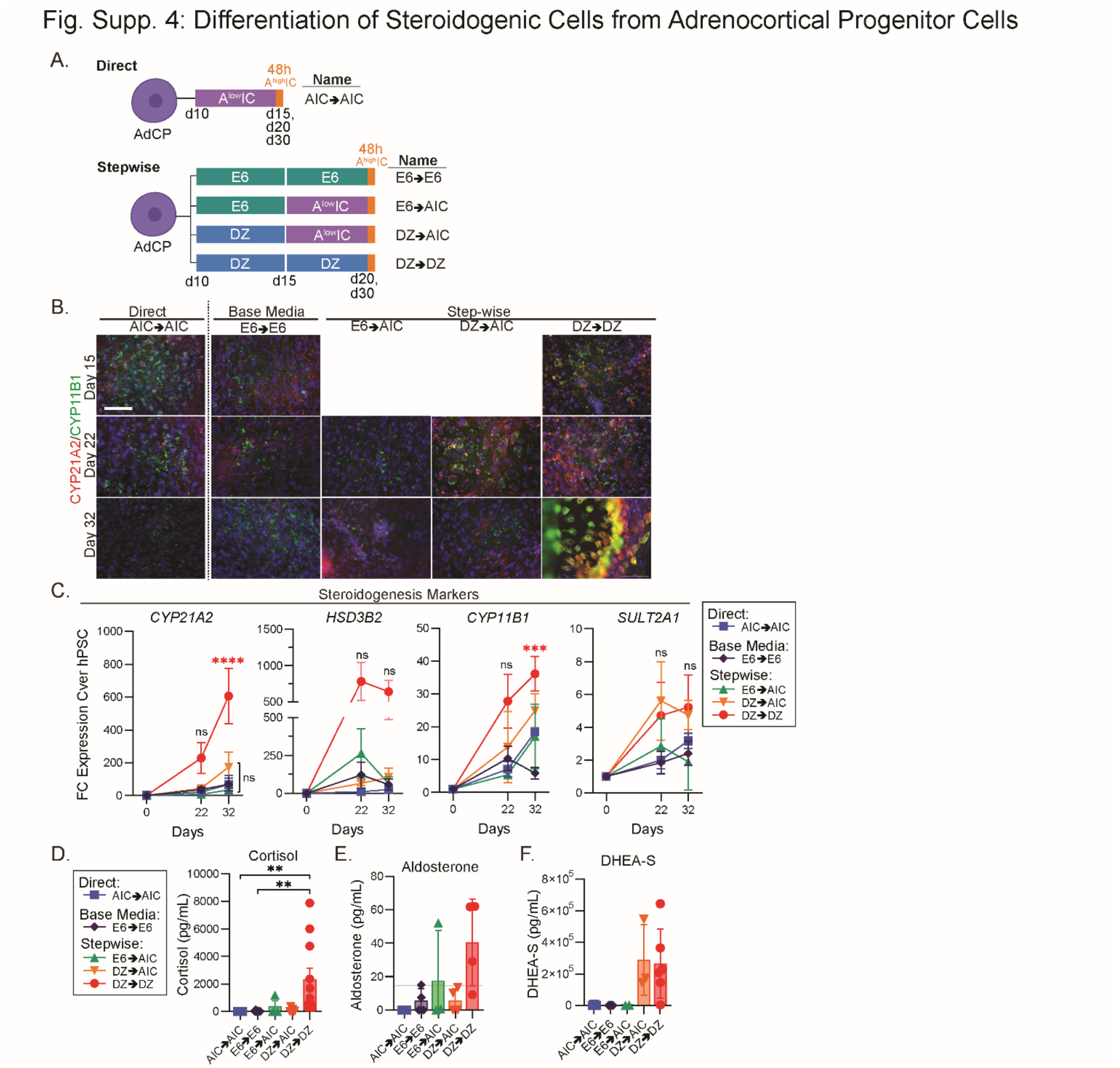


**Fig. Supp. 4: Differentiation of Steroidogenic Cells from Adrenocortical Progenitor Cells**
(**A**) Diagram of approaches to generate steroidogenic cells from AdCPs. (**B**) IF of CYP21A2 and CYP11B1 with nuclear counterstain DAPI (blue) at days 15, 22, and 32 of differentiation (**C**) Steroidogenesis markers for each zone over time (RT-qPCR; n = 2-6). Two-way ANOVA with Tukey's multiple comparisons correction. All comparisons depicted are between each condition versus base media (E6→E6). Asterisk color indicates significant condition. (*CYP21A2*: ****p <0.0001; *CYP11B1*: E6→E6 versus DZ→DZ, ***p= 0.0001). All other significance results reported in raw data file uploaded to Mendeley. (**D**) Cortisol release from cells differentiated with each approach in (**A**) upon 48 hours of stimulation on day 30 (ELISA; n=4-9; AIC→AIC versus DZ→DZ, **p = 0.005; E6→E6 versus DZ→DZ, **p=0.004; all others n.s.). Wallis test (non-parametric one-way ANOVA, Dunn's multiple comparison correction). (**E**) Aldosterone secretion in each condition. (**F**) DHEA-S secretion in each condition after 48 hours of AIC stimulation on day 30 (ELISA; n = 3-5, n.s.). Kruskal Wallis test. All graphs in this figure show mean ± SEM. Abbreviations: AdCP = adrenocortical progenitor cells; AIC = ACTH+IGF2+CRH; DZ = definitive zone; E6 = base media.

Fig. Supp. 5 - Characterization of Response to ACTH Stimulation, Extended Data

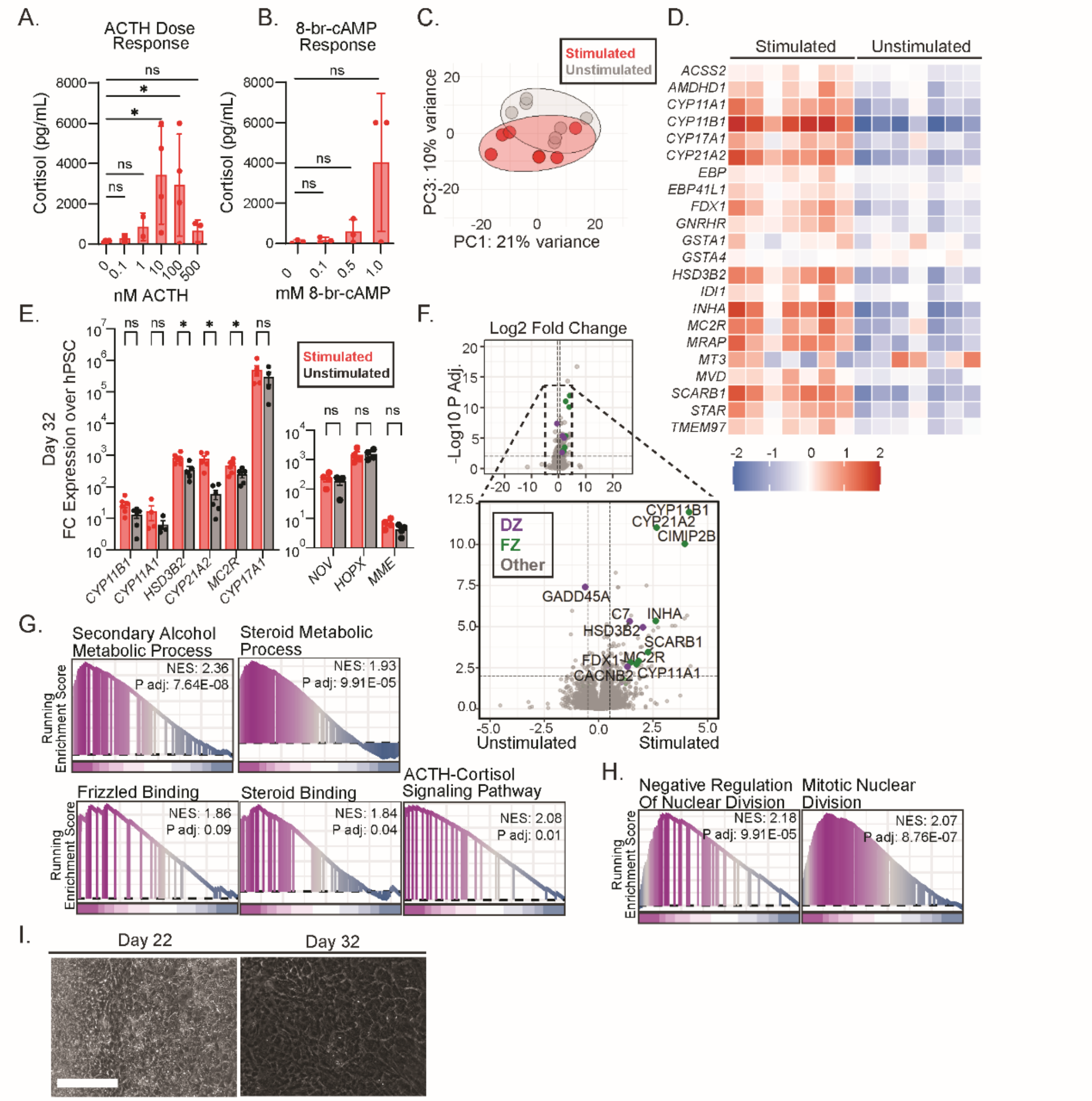


**Fig. Supp. 5: Characterization of Response to ACTH Stimulation, Extended Data**
(**A**) Cortisol secretion in day 32 matured DZ (mDZ) cells was measured by ELISA across a range of ACTH concentrations ranging from 0.1nM to 500nM. One-way ANOVA with uncorrected Fisher's LSD. (0 vs 10nM, *p=0.0178; 0 vs 100nM *p=0.0382). (**B**) Cortisol secretion was also measured in response to 48-hour treatment of day 30 mDZ cells with 0.1, 0.5, and 1.0 mM 8-br-cAMP. One-way ANOVA and Tukey's multiple comparisons test (n.s.). (**C**) PCA plot of PC1 and PC3 of stimulated and unstimulated samples analyzed by bulk RNAseq. (**D**) Heatmap of z-scored expression of ACTH-response transcripts in each stimulated versus unstimulated mDZ sample. Data corresponds to the averaged values shown in **Fig. 3F**. (**E**) RT-qPCR for steroidogenesis markers (left) and DZ markers (right) in stimulated and unstimulated mDZ at day 32. Ratio paired t-tests with Holm-Šídák method multiple comparison correction (*HSD3B2*: *p adj.=0.0166; *CYP21A2*: *p adj.=0.0296; *MC2R*: *p adj.=0.0296). (**F**) Volcano plot of differential expression results for stimulated (positive log2 fold change, right side of plot) vs unstimulated (negative log2 fold change, left side of plot) mDZ. Highlighted are DZ markers or FZ markers. (**G**) Leading edge plots for select GSEA results related to steroidogenesis and cell signaling. (**H**) Leading edge plots for select cell cycle terms. Leading edge plots relate to GSEA results in **Fig. 3I**, **J** and depict thoroughness of GSEA term enrichment based on differential gene expression list. (**I**) Phase contrast microscopy of mDZ culture at days 22 and 32 of differentiation. Scale = 100μm. All graphs in this figure show mean ± SEM. Abbreviations: GSEA = gene set enrichment analysis; mDZ = matured DZ; NES = net enrichment score; PCA = principal component analysis.

Figure Supp. 6: All Identified Consensus Zonation Markers

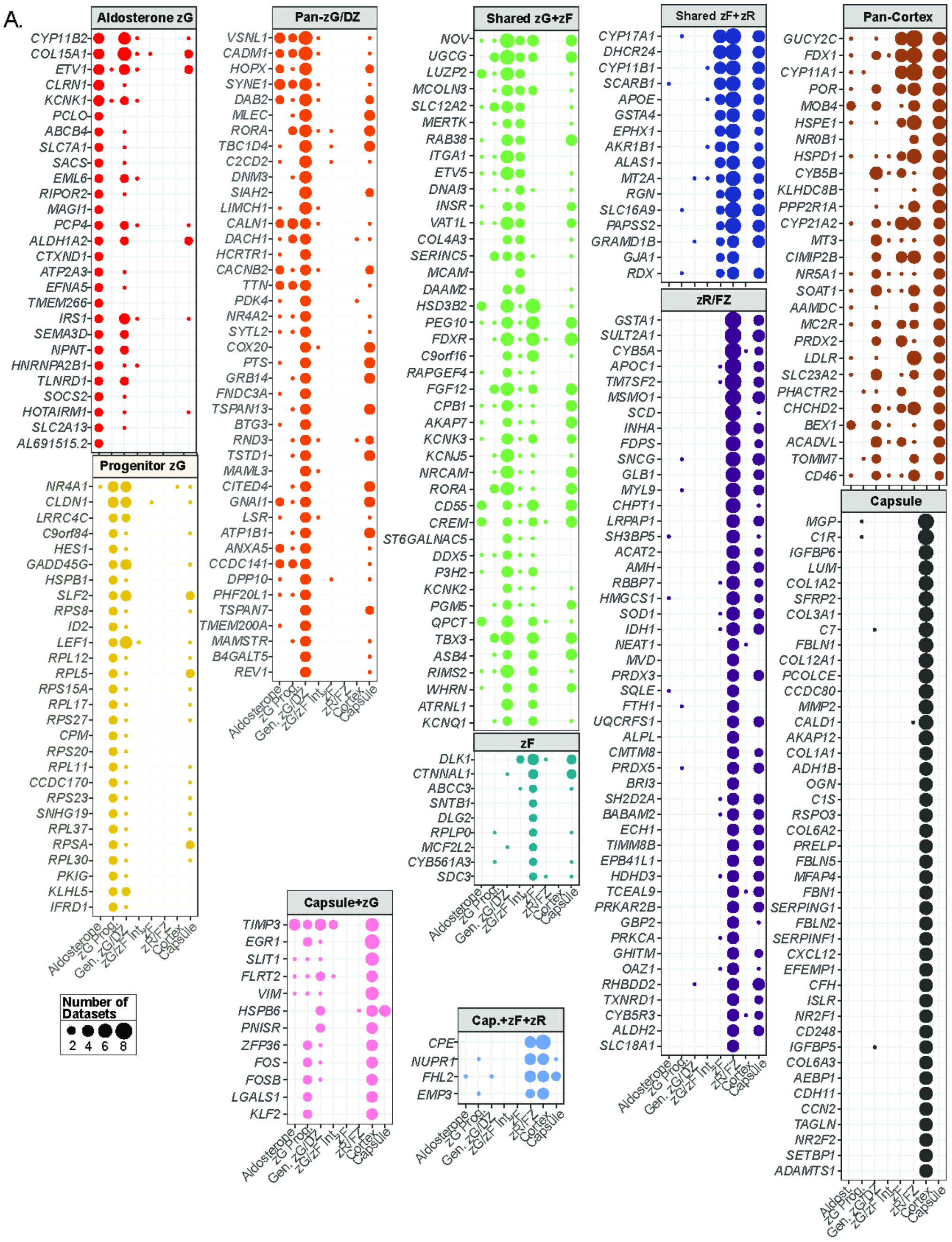


**Fig. Supp. 6: All Identified Consensus Zonation Markers**
(**A**) Dotplot of consensus markers of adrenal capsule and cortex cell types containing an expanded dataset corresponding to **Fig. 4C**. Dot size corresponds to the number of times each marker (y-axis) was present in a cell type cluster from the original publication (x-axis). All markers in **Fig. 4C** are present in this figure along with all other markers identified to be specific but that were not in the top 15 genes listed.
Abbreviations: DZ = definitive zone; FZ = fetal zone; zG = zona glomerulosa; zF = zona fasciculata; zR = zona reticularis.

### Figure Supp. 7: Temporal Marker Specificity Extended Data

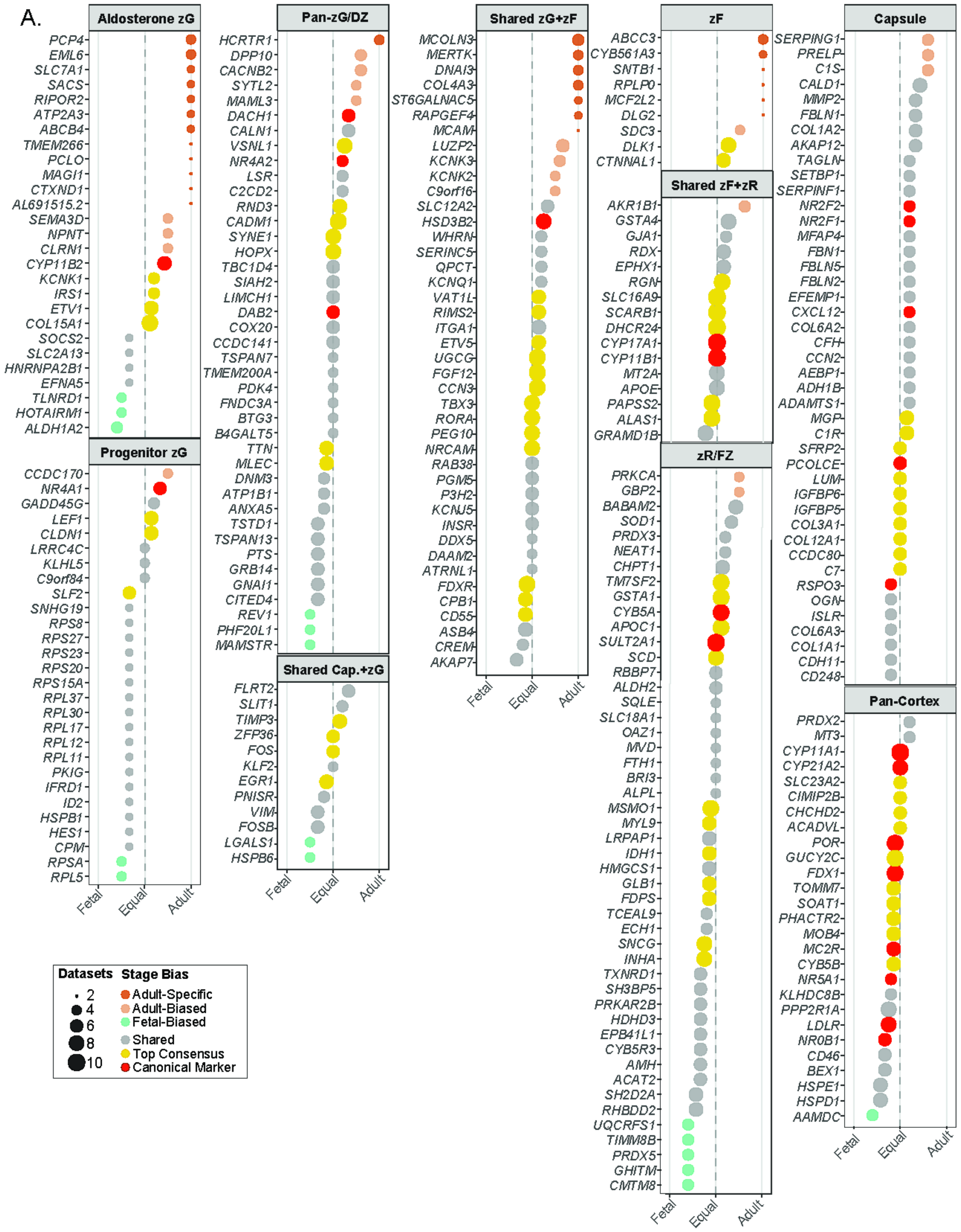


**Fig. Supp. 7: Temporal Marker Specificity Extended Data**
(**A**) Diverging dotplot containing an expanded dataset from **Fig. 4E**. Genes with fetal specificity (left of center), strongest adult specificity (right of center), conserved expression across life stages (closest to center). Established/canonical cell type markers and strongest conserved markers (closest to center combined with the largest number of datasets the gene is differentially expressed in this cell type cell type) are each highlighted. Distance from center line indicates higher counts in one life stage category versus the other. Size of dot represents number of times a gene was present in the cell type category the in the original datasets. Abbreviations: DZ = definitive zone; FZ = fetal zone; zG = zona glomerulosa; zF = zona fasciculata; zR = zona reticularis.
